# Glycocalyx-mediated Cell Adhesion and Migration

**DOI:** 10.1101/2020.06.12.149096

**Authors:** Samuel Schmidt, Bettina Weigelin, Joost te Riet, Veronika te Boekhorst, Mariska te Lindert, Mietske Wijers-Rouw, Barbara Lelli, Lorenz Rognoni, Marina Krause-Vortmeyer, Anthea Messent, Luisa Bracci, Kay-Eberhard Gottschalk, Stephan Kissler, Martin J. Humphries, Dirk J. Lefeber, Jack Fransen, Peter Friedl

## Abstract

Cell migration is a force-dependent adaptive process mediated by integrin-dependent adhesion as well as other yet poorly defined interactions to the extracellular matrix. Using enzymatic multi-targeted digestion of sugar moieties on the surface of mesenchymal cells and leukocytes after interference with integrin function, we demonstrate that the surface glycocalyx represents an independent adhesion system. The glycocalyx mediates cell attachment to ECM ligand in the 100-500 pN force range and amoeboid migration in 3D environments *in vitro* and *in vivo*. Glycan-based adhesions consist of actin-rich membrane deformations and appositions associated with bleb-like and other protrusions forming complex-shaped sub-micron contact sites to ECM fibrils. These data implicate the glycocalyx in mediating generic stickiness to support nanoscale interactions (nanogrips) between the cell surface and ECM, mechano-coupling, and migration.

## Introduction

Cell shape, polarity and anchorage, as well as migration across surfaces depend upon the function of integrin adhesion receptors, which form transient focalized actin-containing adhesion complexes that control cytoskeletal organization, mechano-transduction and intracellular signaling^1,2^. In cancer metastasis, integrins mediate cell invasion and represent candidate targets for pharmacological interference^3-5^. In contrast to 2D migration models, interference with β1 integrin function in 3D environments, where cells migrate through substrate of complex geometry and along confining interfaces, provides only incomplete or no inhibition of migration^6-9^. The mechanisms mediating cell-substrate coupling and migration when integrin availability is low or absent, mediated by “friction”^9,10^, physical intercalation^11^, or alternative adhesion systems^12^ remains elusive. Our aim was therefore to identify the cellular and molecular mechanisms of integrin-independent cell-matrix interaction, force generation, and migration within collagen-rich interstitial tissue *in vitro* and *in vivo* when integrin-mediated adhesion is marginalized or absent.

## Results

### Mesenchymal-to-amoeboid transition after interference with integrins

Fibrillar collagen is the predominant extracellular matrix (ECM) structure in mammalian tissues and recognized with high affinity by integrins *α*1β1, *α*2β1, and *α*11β1, and weakly by *α*Vβ3^13^. Invasive MV3 melanoma cells, which express β1 and β3, but lack all other integrin β-chains (Extended Data Fig. 1a), utilize *α*2β1 integrins for efficient migration within in vitro reconstituted 3D collagen matrix^14^ and collagen remodeling^15^. During migration through 3D fibrillar collagen, up to 90% of MV3 cells adopted an elongated, spindle-shaped morphology and migrated at speeds similar to the movement of mesenchymal cells *in situ*^16^ (Fig. 1a). Graded interference with β1 integrins was induced either by adhesion-perturbing mAb 4B4 at concentrations saturating the epitope up to 90% (Extended Data Fig. 1a and 1c) or by stable shRNA-based β1/β3 integrin downregulation (Extended Data Fig. 1d) combined with additional anti-β1 antibody based interference which achieved >99% epitope reduction for β1 without detectable β3 integrin at the cell surface (Extended Data Fig. 1e, f). Individual- and dual-integrin interference strategies reduced migration speed by >80% but failed to achieve immobilization with residual slow migration speed of 0.02-0.15 µm/min (Fig. 1a). As consequence of interference with integrins, the spindle-shaped, elongated morphology converted to an ellipsoid cell shape with multiple dynamic blebs and occasional filopodia in contact with collagen fibrils and focalizations of β1 integrin and filamentous actin at contact sites to collagen fibers converted to diffuse distribution (Fig. 1b, arrowheads). These data show for metastatic melanoma cells a transition from mesenchymal to amoeboid migration when integrin availability is limited.

**Fig. 1.**
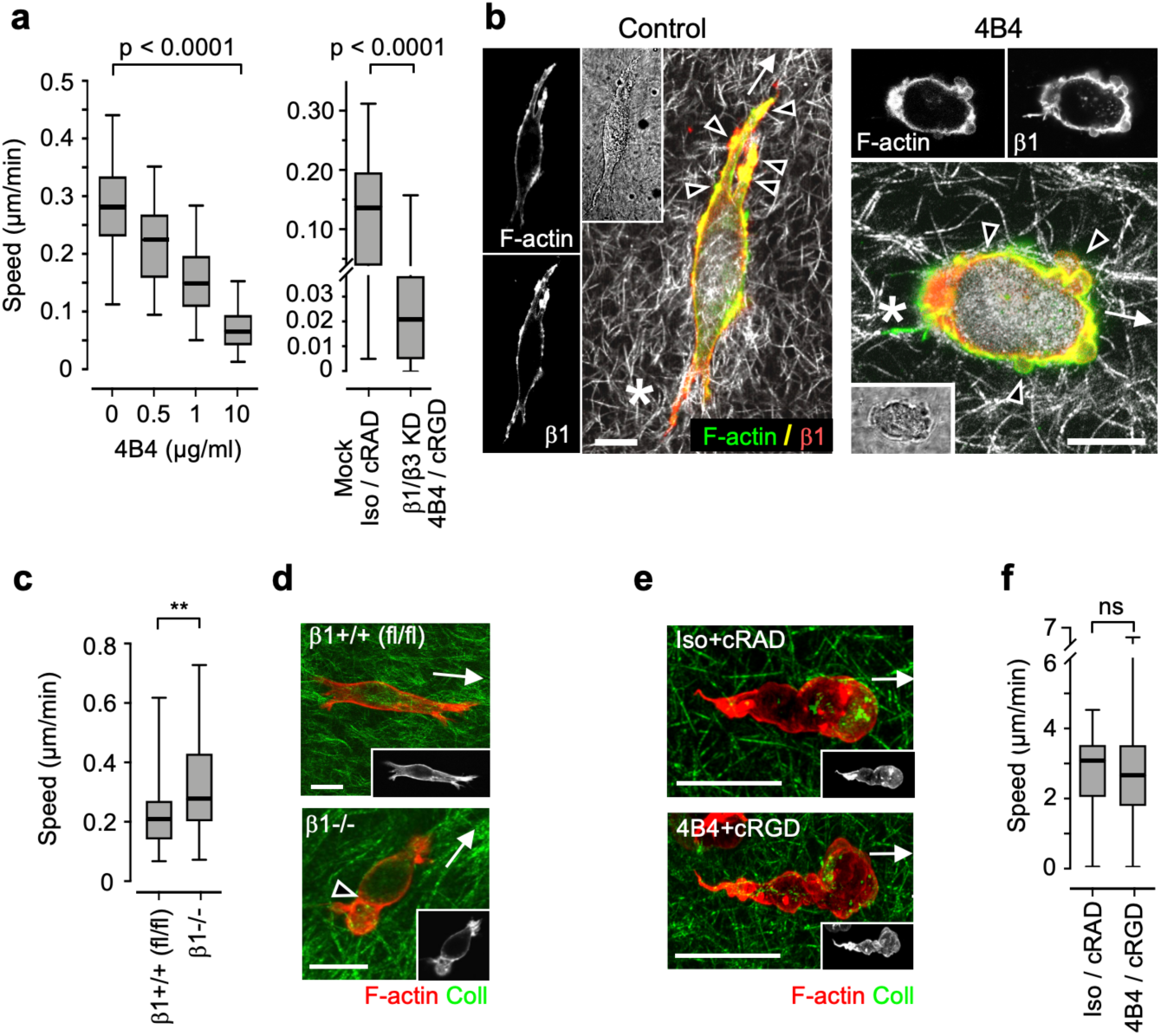
Conversion to amoeboid migration in mesenchymal tumor cells, fibroblasts and T lymphoma cells after interference with integrins. (a) Speed change in MV3 cells by mAb 4B4 or β1/β3 shRNA combined with mAb 4B4 and cRGD. Median population speed (single cell tracking, observation period 24 h, left panel and 6h, right panel, 38-117 cells, 3 independent experiments). (b) Cell morphology, F-actin and β1 integrin distribution (arrowheads) for control and mAb 4B4 treated cells. Arrows, direction of migration, based on retraction fibers from the cell rear (asterisks). Bars, 10 µm. (c) Median migration speed (23-67 cells, 3 independent experiments) and (d) morphology of wild-type and β1-/-MEFs. F-actin (red; grey in insets), collagen fibers (green; reflection signal), and constriction rings (arrowheads). Bars, 100 µm. (e) Morphology and (f) median migration speed (73-75 cells, 3 independent experiments) of human Molt-4 cells in the absence or presence of mAb 4B4 and cRGD. F-actin (red; grey in insets), collagen fibers (green; reflection signal) and direction of migration (arrows). Bars, 20 µm. (a) P values, Kruskal-Wallis with Bonferroni post-test and (c, f) Non-paired Mann-Whitney test, 2 tailed. Box and whisker plots show 25-75 percentiles (box), the median (middle line) and 5/95 percentiles (whiskers). See also **Extended Data Fig. 1-3**.

To test whether β1 integrins are dispensable for cell migration in non-cancer cells, β1-deficient murine embryonic fibroblasts (MEFs) were tested (Extended Data Fig. 2a-c). β1-/-MEFs employed a rounded amoeboid migration type with increased speed compared to β1+/+(fl/fl) MEFs (Fig. 1c, d) reaching 0.16-0.28 µm/min. With reduced cell elongation (Exdended Data Fig. 2d), directional persistence was not compromized compared to β1+/+(fl/fl) MEFs (Extended Data Fig. 2e, Supplementary Video 1) but, given the increased speed, integrin deficiency did not compromise effective migration from multicellular spheroids (Extended Data Fig. 2f).

**Fig. 2.**
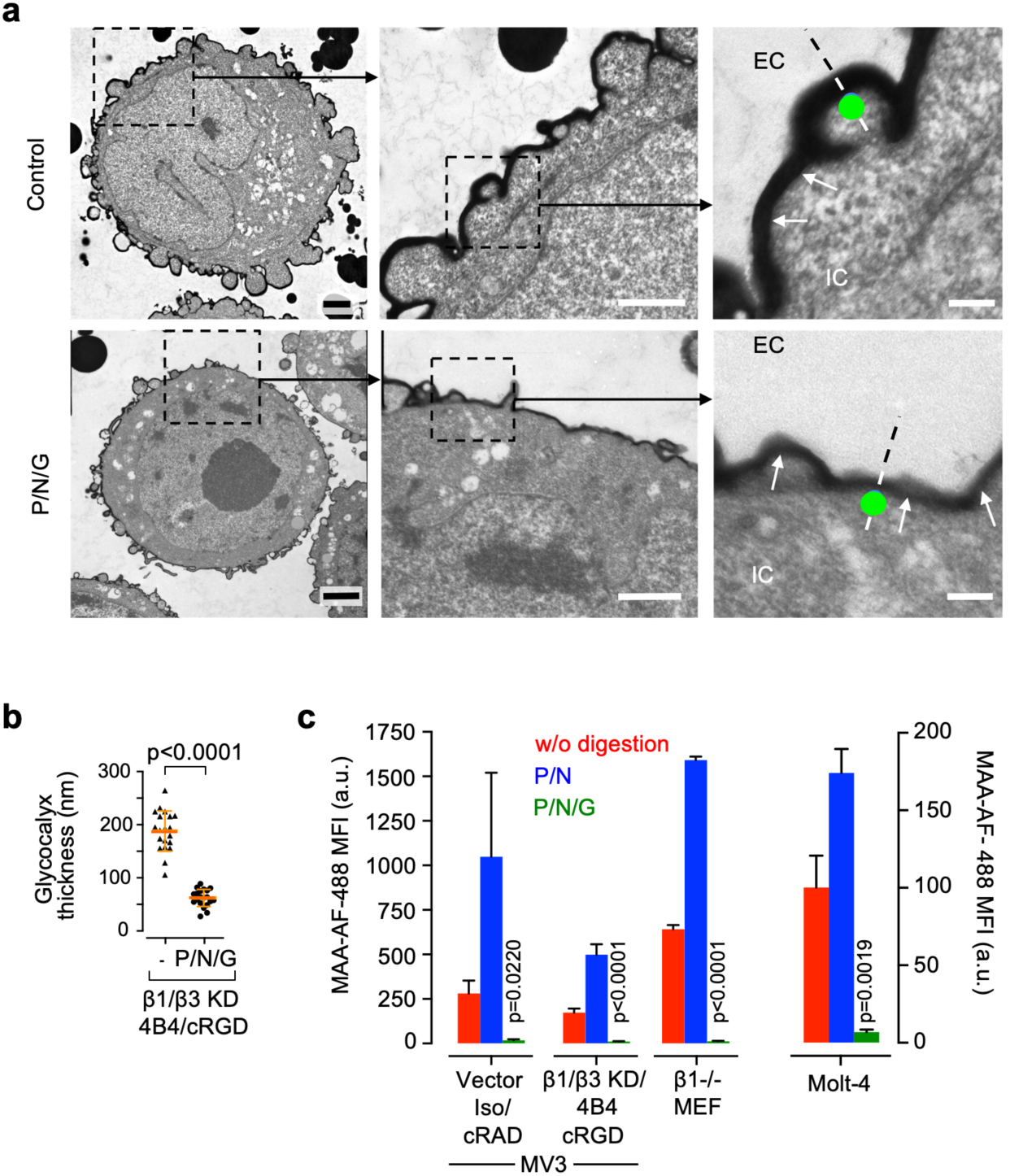
Enzymatic removal of the glycocalyx. (a, b) Ultrastructure of the glycocalyx. Glycocalyx thickness analysis by transmission electron microscopy (a) and quantitative image densitometry (b). MV3 β1/β3KD cells before and after glycan removal with P/N/G glycosidase cocktail followed by glycan detection with Ruthenium red staining. Blue dot in (a), plasma membrane. Dashed lines, cross-sections perpendicular to the cell surface used for image analysis in (b). Bars, 1 µm (left), 1 µm (middle), 200 nm (right). (b) Representation of the means and SD (10 cells, 3 independent experiments), derived from densitometry profiles shown in Fig. S4B. (c) Digestion efficacy of surface glycans on MV3, β1-/-MEF and Molt-4 cells. β1-4 galactose was detected with Maackia amurensis agglutinin (MAA) and fluorescence intensity was assessed by flow cytometry. Means and SEM from 3 (MV3 vector), 7 (MV3 β1/β3KD) and 3 (MEF and Molt-4) independent experiments. P values (all graphs), (b) non-paired Mann-Whitney test, 2 tailed and (c) non-paired t-test, 2 tailed. See also **Extended Data Fig. 4**.

Together, these findings support the concept that in mammalian mesenchymal cells the availability of integrins determines the motility programs^17^. Mesenchymal-like migration of fibroblasts and some cancer cell types results from elongated spindle-shaped morphology, focalized adhesion sites, and high traction force towards the ECM whereas low integrin availability generates weak mechanotransduction by means of poorly focalized adhesion sites and actin cytoskeleton, similar to leukocytes and otherwise adhesive cells moving in non-adhesive 3D environments^9,10^.

As confirmatory model, integrin-expressing (Extended Data Fig. 3a, b) but integrin-independent migrating Molt-4 human T lymphoma cells developed amoeboid, rounded shape with an elongated uropod, migration speed between 2 to 4 µm/min and directional persistence in both untreated control conditions or in the presence of mAb 4B4 and cRGD interfering with β1 and *α*Vβ3 integrins, respectively (Fig. 1e, f, Extended Data Fig. 3c, d). These results confirm for a 3D ECM model that moving cells possess integrin-independent interaction mechanisms with fibrillar collagen to maintain migration.

**Fig. 3.**
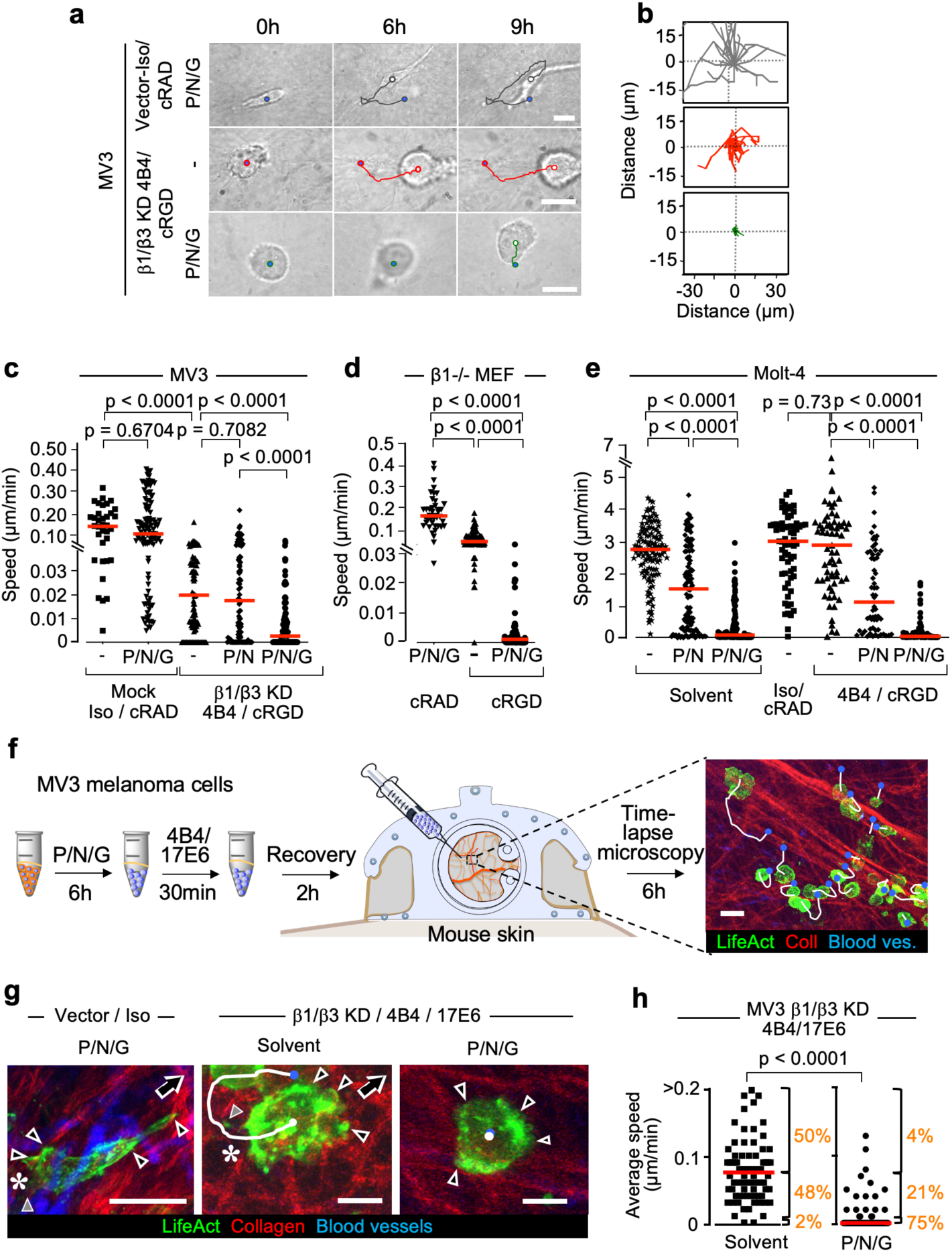
Surface-glycan dependent cell migration in vitro and in vivo. (a-e) Effects of enzymatic surface-glycan removal on cell migration in 3D collagen lattices. Migration rates in 3D collagen matrices of MV3 melanoma cells (a-c), murine β1-/-embryonic fibroblasts (d) and Molt-4 cells (e) after treatment with glycosidases P/N or P/N/G. Time-lapse sequences (a; bright-field microscopy), path organization and single-cell speed distribution or the indicated conditions (c; 38-117, d; 41-83, e; 60-111 cells, 3 independent experiments). Red lines, medians. (f-h) Impact of surface glycan removal on MV3 cell migration in vivo. (f) Workflow of orthotopic injection into the collagen-rich deep mouse dermis followed by intravital time-lapse microscopy of Lifeact-expressing MV3 vector control and MV3 β1/β3KD cells without or after glycan removal. Bar, 20 µm. (g) Representative zooms of cell morphology and migration paths (white lines) over 8-9h time-lapse period. Arrows, direction of migration. Grey arrowhead, microparticle released from the cell rear (Asterisk). Bars, 20 (Vector/Iso) and 10 µm (β1/β3KD). (h) Migration speed of MV3 β1/β3KD cells additionally treated with mAb 4B4 and 17E6 without or after treatment with P/N/G glycosidases (78 cells/condition, 3 independent mice). (c-e, h) Red lines, medians. P values (all graphs), non-paired Mann-Whitney test, 2 tailed. See also **Extended Data Fig. 4 and 5**.

We explored alternative collagen receptors^12^ that might compensate the loss of integrin-mediated adhesion and migration in MV3 cells, including syndecan-1, discoidin domain receptors (DDR) 1 and 2, and proteoglycan CD44. DDR-1 and -2 or syndecan-1 were not expressed by MV3 cells after culture in 3D collagen matrix and interference with integrins (Extended Data Fig. 3e, f). CD44 was expressed, however CD44 perturbing antibody Hermes-1 did not affect the migration of MV3 cells after β1/ β3 integrin knockdown in 3D fibrillar collagen (Extended Data Fig. 3g, h). These data argue against an important role of these receptors in mediating amoeboid movement in MV3 cells.

### Multi-enzyme removal of surface glycans

Besides cell surface receptors, the surface glycocalyx can interact with proteins and other materials, through carbohydrate-binding domains^18^ or unclassified ionic and non-ionic bonds providing generic stickiness^19^. We therefore hypothesized that non-specific low-affinity interaction of mammalian cells with collagen fibers could be mediated by the glycocalyx^20-22^ and/or even non-adhesive 3D cell intercalation^11,23^.

In mammalian cells, the glycocalyx forms a thick, polar layer, which is composed of various classes of glycoconjugates, that are directly or indirectly coupled to the cell membrane^24^. These molecules include glycosaminoglycans (GAGs) chains of cell surface proteoglycans including heparan sulfate and chondroitin sulfate and glycoproteins and glycolipids containing N- and O-linked glycoconjugates (Extended Data Fig. 4a)^25^. Highly sulfated GAG side chains of proteoglycans are known to interact with most ECM components, including fibrillar collagen and fibronectin^26^. Depending on the cell type, the glycocalyx may project hundreds of nanometers to several micrometers^4,27^, and transmission electron microscopy analysis of MV3 cells demonstrated that these cells expressed a thick glycan layer at the cell surface of approximately 200 nm in thickness (Fig. 2a, b, Extended Data Fig. 4b).

**Fig. 4.**
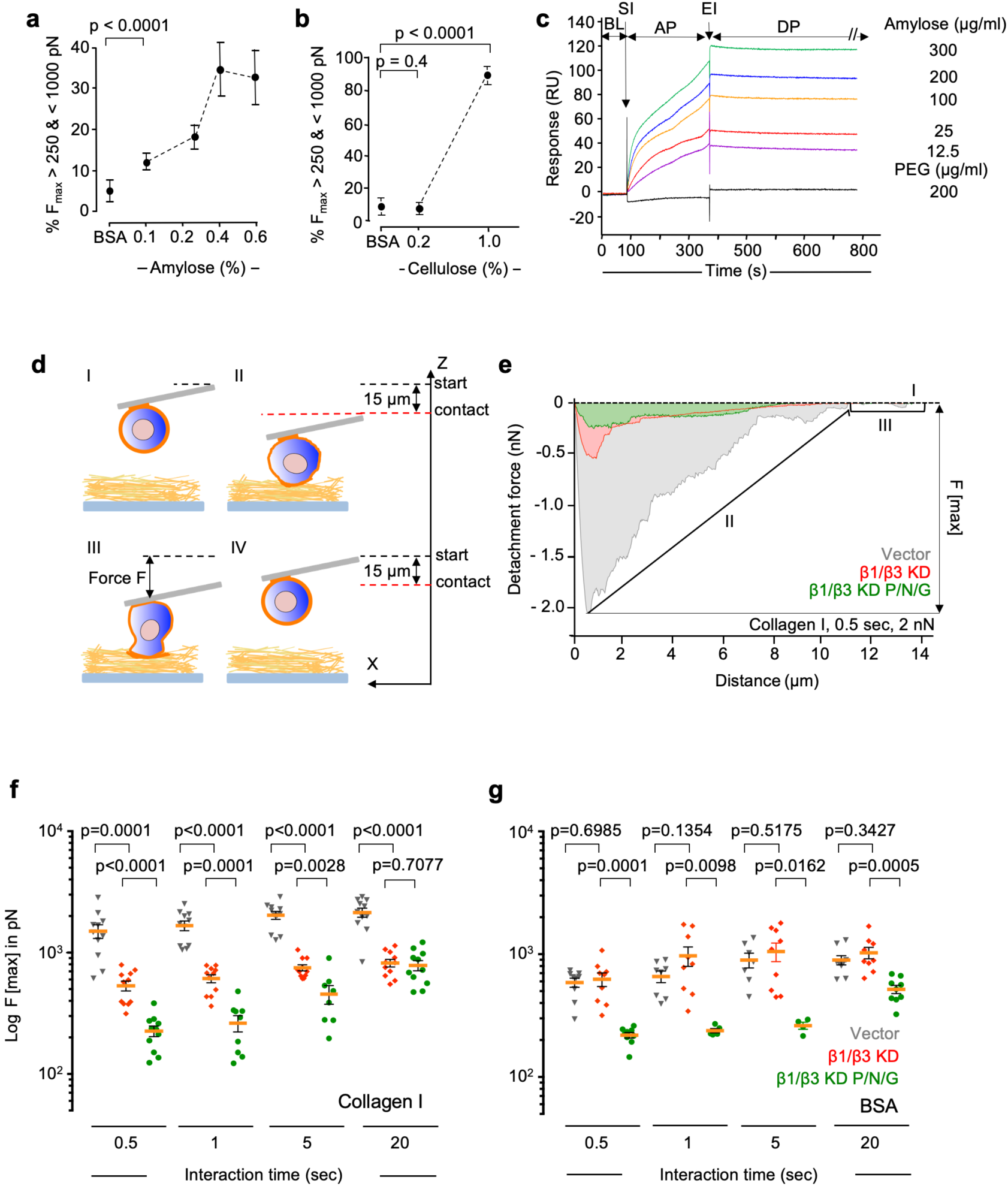
Probing of glycan-binding to collagen fibers and studying the impact of enzymatic digestion of the glycocalyx on glycan-mediated cell binding to collagen. Atomic force spectroscopy of glycan-binding to collagen fibers. (a) Representation of ‘high-force’ interactions above 250 pN with increased coating concentration of amylose. (b) Force distribution for low- and high-concentration of cellulose. Means and SEM of 1,600 to 10,000 force curves per sample from 3 independent experiments. P values, non-paired Mann-Whitney Wilcoxon test, 2 tailed. (c) Glycan-mediated binding affinities to collagen fibers monitored by surface plasmon resonance (SPR). Overlay of amylose-collagen I affinity sensorgrams obtained by measurements of changes in the SPR response of increased amylose concentrations. 1 representative sensorgram out of 2 independent experiments for each amylose concentration and the PEG control. (d-g) Atomic force-based life cell spectroscopy of cell-surface glycan-binding to collagen. (d) Coupling of a single cell to AFM cantilever to probe the force required for cell detachment F [max] from a 3D fibrillar collagen surface. (e-g) Live-cell atomic force spectroscopy of untreated MV3 control cells, MV3 β1/β3KD cells and MV3 β1/β3KD cells additionally treated with P/N/G glycosidases. (e) Force-distance curve of a single force measurement cycle between MV3 cell and collagen surface, including start position (I), retraction (II), and detachment phase (III). The maximum detachment force exerted on the cantilever F [max] was calculated from the peak to background level. (f, g) Interaction forces of MV3 cells to fibrillar collagen I (f) and BSA-coated surface (g) after 0.5 s, 1 s, 5 s or 20 s interaction time. Pooled values of maximum retraction forces to a mean value shown as Log F [max] of 5 individual measurements per cell (orange lines show means with SEM). P values (all graphs), non-paired t-test, 2 tailed. See also **Extended Data Fig. 6 and 7**.

To address the role of surface glycans in cell migration, protein- and lipid-linked glycoconjugates were enzymatically removed from the surface of live cells by a two-step glycosidase treatment. The enzymatic digestion combined hyaluronidase, heparitinase, chondroitinase and neuraminidase with galactosidase, the latter hydrolyzing β-1,4 coupled galactose residues in N- and O-linked glycans and glycolipids, lowered cell surface heparan- and chondroitin sulfate by ∼98% and dermatan sulfate by ∼95% (Extended Data Fig. 4 c). When combined with sialic acid removal by neuraminidase, this treatment resulted in reduction of sialic acid residues by ∼80% in MV3 control and β1/β3KD cells, by ∼95% in Molt-4 cells (Extended Data Fig. 4c) and an exposure of subjacent β-1,4 coupled galactose residues (Fig. 2c, Extended Data Fig. 4d; step 1 “P/N”). Additional treatment with β1-4 galactosidase strongly reduced the glycocalyx thickness compared to untreated control cells (Fig. 2b) and removed β-1,4 coupled surface galactose by >90% in MV3 control and β1/β3KD cells, >98% in MEF β1-/- and >94% in Molt-4 cells (Extended Data Fig. 4d; step 2 “P/N/G”) without negatively impacting cell viability (Extended Data Fig. 4e). The loss of cell surface β-1,4 galactose was maintained for at least 6-9h (MV3), 24-92 h (MEF β1-/-) and 5-6 h (Molt-4) followed by a stepwise recovery (Extended Data Fig. 4f, g). Likewise, during the recovery, the cells showed consistent viability (Extended Data Fig. 4h).

To verify that the digestion procedure was non-toxic and further did not compromise the basic migration ability through integrins, MV3 control cells expressing β1/β3 integrin after enzymatic digestion (P/N/G) together with non-inhibitory cRAD showed normal baseline migration and mesenchymal phenotype (Fig. 3a-c). Likewise, β1-/-but β3 integrin-expressing MEFs developed similar morphological phenotypes and unperturbed migration speeds after enzymatic digestion compared with β1+/+(fl/fl) MEFs (Fig. 3d versus 1c). These data indicate unperturbed migration and, hence, cell viability and cytoskeletal activity after glycosidase treatment.

### Surface-glycan dependent cell migration

While mesenchymal migration of integrin-competent control cells was unaffected, combining glycan removal and interference with integrin expression severely compromised cell migration and persistence in 3D collagen lattices across cell models. Glycan removal immobilized all cells, despite ongoing cytoskeletal activity (“running on the spot”), irrespective of whether integrin-independent baseline migration was slow in MV3 (Fig. 3a-c, Supplementary Video 2, 3), intermediate in β1-/-MEFs (Fig. 3d, Supplementary Video 4) or fast in Molt-4 cells (Fig. 3e, Supplementary Video 5). Glycan removal was further accompanied by compromised directional persistence (Extended Data Fig. 5b-d) and cell elongation (Extended Data Fig. 5e, f). Cells remained immobile for time periods of 6h (MV3), 24h (MEF) or 5h in Molt-4 cells, after which they gradually regained migration ability (Extended Data Fig. 5a) and β-1,4 galactose on the cell surface (Extended Data Fig. 4f, g). Notably, the enzymatic removal of GAGs and neuraminic acids alone (P/N) did not influence migration of MV3 β1/β3KD cells and β1-/-MEFs (Fig. 3c), whereas the migration of Molt-4 cells was partly compromised (Fig. 3e, P/N). This indicates cell-type specific use of glycan subtypes for maintaining migration^28^.

**Fig. 5.**
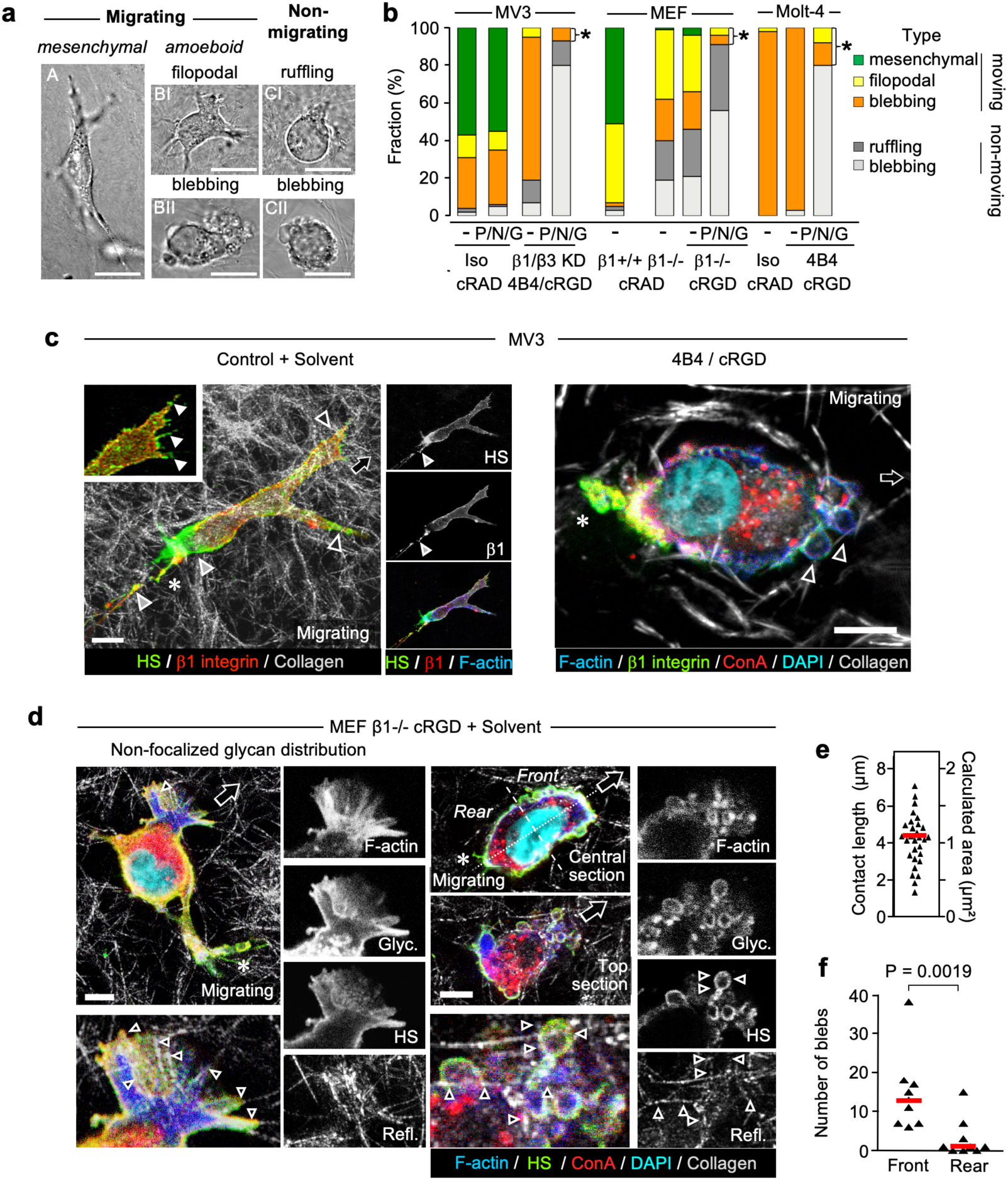
Cell morphology and cellular interactions to collagen fibers of MV3, MEF and Molt-4 cells and the impact of glycan removal. (a) Phenotypes in migrating and non-migrating cells according to shape and polarity after 2 of culture in 3D collagen lattices. (b) Occurrence of MV3, MEF and Molt-4 phenotypes after interference with integrins and surface glycans from 97 (MV3), 100 (MEF) and 92 (Molt-4) cells (2 independent experiments). Asterisks, amoeboid-rounded subsets. Bars, 20 µm. (c-g) Structure and molecular composition of cellular interactions with collagen fibers. MV3 (c) and β1-/-MEF (d) cells were fixed after 2h-culture in 3D collagen lattices. (c, left panel) Spindle-like shape of MV3 cells and localization of heparan sulphate (HS) at interaction sites with collagen fibers (black arrowheads) during migration. HS is accumulated in anterior ruffles and filopodia (inset of individual channels, white arrowheads) and the trailing edge (gray arrowheads). Asterisk, deposited material from the cell rear, confirming the migratory state. Bar, 10 μm. (c, right panel) Spherical, slowly migrating MV3 cell after interference with integrins (mAb 4B4 and cRGD). In fixed cells, concanavalin A stains both plasma membrane and intracellular vesicles. Arrowheads, location of collagen fibers, based on the single-channel reflection images. Arrows, direction of migration, based on retraction fibers or small particles released from the cell rear (asterisks). 1 representative cell for each condition from more than 10 cells, analysed in 2 independent experiments. Bars, 5 μm. (d, left panel) Non-focalized distribution of glycans (stained by concanavalin A) and heparan sulphate (HS) at the surface of actin-rich filopodia of β1-/-MEF cells extending along collagen fibers. 1 representative cell from 4 cells, analysed in 2 independent experiments. (d, right panel) Distribution of glycans and HS on actin-rich cell blebs in contact with collagen fibers. Central section showing surface ‘roughness’ and tangential top section detecting protruding blebs that intercalate between collagen fibers. 1 representative cell from 10 cells, analysed in 2 independent experiments. Bars, 5 μm. (e) Length and calculated area of the interface between filopodia of β1-/-MEFs and collagen fibers. 1 representative experiment with 28 data points from 10 cells, pooled from 2 independent experiments. (f) Number of blebs from central sections in the front and rear half of 9 analysed cells from 2 experiments, according to the dashed lines in (d, D). (e-g) Red lines, medians. P value, non-paired Mann-Whitney test, 2 tailed.

We next sought to address whether the glycan-dependence of cell migration is relevant in collagen-rich interstitial tissue *in vivo*. MV3 β1/β3KD cells were additionally pretreated with integrin-blocking mAbs 4B4 and 17E6, injected into the deep dermis of nude mice and monitored by intravital microscopy for up to 6h (Fig. 3f). Whereas MV3 vector control cells developed spindle-shaped morphology (Fig. 3g, Extended Data Fig. 5g, A, B), integrin targeting caused rounded morphology (Fig. 3g, Extended Data Fig. 5g, C, D) and blebbing movement *in vivo* (Fig. 3h, Supplementray Movie 6). These data indicate that lowering integrin availability in mesenchymal cells results in amoeboid movement irrespective of 3D environments *in vitro*, including microfluidic channels^29^, fibrillar collagen^8^ as well as interstitial tissue *in vivo*. However, when surface glycans were removed before injection into the mouse dermis, MV3 β1/β3KD cells underwent near-complete migration arrest (Fig. 3g, h), but maintained oscillatory shape change as an indication for unperturbed viability (Extended Data Fig. 5g, E, F, Supplementary Video 6). Thus, an intact surface glycocalyx is required to maintain amoeboid migration in collagen-rich tissue *in vitro* and *in vivo* when integrin functions are perturbed.

### Glycan-mediated attachment forces in the pN range

To address directly whether the glycocalyx functions as adhesion system, atomic force spectroscopy (AFS) was used to probe the binding of synthesized glycan polymers and live cells to fibrillar collagen. The tip of the cantilever was functionalized with simple-structured amylose and cellulose that lack modifications such as sulfation or acetylation and the forces to disrupt the bonds upon cantilever retraction were recorded (Extended Data Fig. 6a). With increasing amylose density on the cantilever, the binding to collagen was dose-dependently strengthened (Fig. 4a, Extended Data Fig. 6b; upper graph). Besides background-level attachments (40-120 pN), amylose or cellulose also enabled stronger bonds with unbinding forces reaching 300-800 pN (Fig. 4a, b, Extended Data Fig. 6b). Effective interaction of amylose with immobilized collagen was confirmed by surface plasmon resonance detection (Extended Data Fig. 6c). After engagement, bonds between amylose and monomeric collagen were stable for >15 min, irrespective of the applied amylose concentration (Fig. 4c, Extended Data Fig. 6d). A wash-out was applied to release amylose from the interaction (Extended Data Fig. 6d), demonstrating reversibility of binding. Thus, multivalent polysaccharides which lack charged side chains effectively adhere to monomeric and fibrillar collagen.

**Fig. 6.**
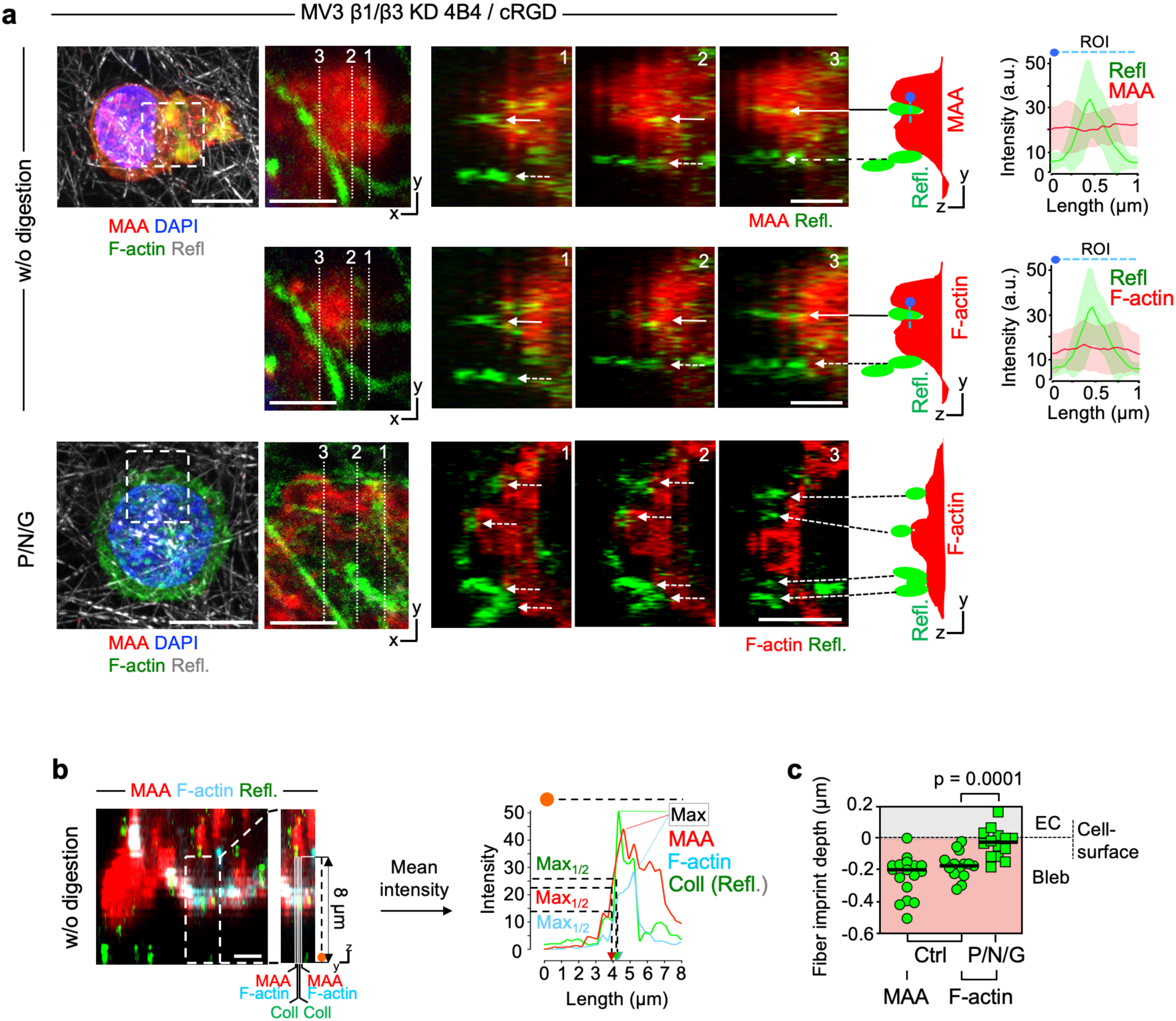
Organization of glycan-dependent cell-matrix interactions. Diffraction-limited confocal microscopy of glycan-dependent interactions with collagen fibers. MV3 β1/β3KD cells were non-treated (a, upper panel w/o digestion, b) or received P/N/G digestion (a, lower panel), embedded in 3D collagen and fixed after 90 min for detection of β-1,4 galactose residues (Maackia amurensis agglutinin, MAA), F-actin and collagen fibers (reflection). Images represent the xy and yz projections from serial z-scans (positions indicated as dotted lines) with relative positions of cross-sectioned fibers (numbered 1-3) and cell surface represented as cartoons. Arrows, collagen fibers (green) located colocalized with MAA and F-actin-enriched protrusions (blebs, red). Dashed arrows, collagen fibers laterally intercalating with the cell surface. 1 representative cell out of 4 cells per condition analyzed in 2 independent experiments. Bars, 10 µm (overview images), 2 µm (ROIs). Mean densitometry curves with SD of MAA or F-actin (red) and collagen fibril intensity (green) based on 20 curves in cross-sectioned bleb-like protrusions (4 MV3 β1/β3KD cells, 2 independent experiments). (b) Quantification the imprint depth of collagen fibers into cell blebs. YZ projection of a MAA (red) and F-actin (cyan) positive membrane bleb. Dashed box, region of interest used for densitometry analysis. White traces indicate the measuring lines with a constant length of 8 µm and a width of 5 pixels. Bar, 2 µm. Representative overlay of intensities alongside the measuring lines in a cross-sectioned MV3 β1/β3KD cell bleb for MAA (red), F-actin (cyan) and collagen I (green) including points of maximal (Max) and half-maximal (Max_1/2_) intensities from 2 independent experiments. (c) Median fiber imprint depth, based on MAA and F-actin signal, as detailed in (b), from 16 cross-sectioned blebs (4 cells/ condition, 2 independent experiments). (c) Black lines, medians. P value, non-paired Mann-Whitney test, 2 tailed. See also **Extended Data Fig. 8**.

To determine binding forces between the glycocalyx and fibrillar collagen by a live-cell strategy, single-cell AFS was applied. Individual MV3 or Molt-4 cells were mounted to the tip of the cantilever and binding forces between cell and substrate were recorded upon cantilever retraction (Fig. 4d, Extended Data Fig. 6e). To probe both fast and slow bond formation with collagen fibrils independently, interaction durations ranging from 0.5 to 60 s (MV3 cells) and 0.5 to 20 s (Molt-4 cells) were applied. Sub-second contacts with defined interaction pressure typically enable rapid adhesion mediated by the cell surface, whereas longer interaction times are expected to additionally engage more complex cytoskeletal remodeling and secondary receptor aggregation^30,31^. In MV3 control cells expressing integrins, short interaction times (0.5 s) enabled variable interaction forces ranging from 600 pN up to 2.8 nN for individually probed cells (Fig. 4e, f, Extended Data Fig. 6f; 0.5 s contact time). In contrast, Molt-4 cells expressing integrins showed a narrow range of interaction forces, from 350 pN up to 1 nN for individually probed cells (Extended Data Fig. 7a; 0.5 s contact time). In MV3 cells stepwise increase in contact time caused elevated peak forces of collagen binding, which was less pronounced in Molt-4 cells (Fig. 4f, Extended Data Fig. 6f, 7a; 20 s contact time). After interference with integrins MV3 β1/β3KD cells were unable to mount very high (>1 nN) but retained moderate peak forces that reached 500-800 pN (Fig. 4e, f, Extended Data Fig. 6f; 0.5 s contact time) and underwent weak adhesion strengthening after 60 s contact time (Extended Data Fig. 6g). In Molt-4 cells peak forces remained constant even after blockage of integrins (Extended Data Fig. 7a). Additional glycosidase treatment in MV3 β1/β3KD cells resulted in a substantial further reduction of peak force, which was even more pronounced in Molt-4 cells (Fig. 4e, f, Extended Data Fig. 6f, 7a; 0.5 s contact time). Diminished adhesion after glycosidase treatment was time-dependent for MV3 β1/β3KD and Molt-4 cells and reverted after 20 s contact time in MV3 β1/β3KD cells (Fig. 4f, Extended Data Fig. 6f, 7a). The time dependence of peak force suggests slow adhesion strengthening, such as by glycan residues very near the plasma membrane which persisted after enzymatic treatment (compare Fig. 2a, b). These data indicate that the glycocalyx supports fast binding (MV3 cells) and slightly prolonged binding (Molt-4 cells) to collagen in the pN range independent of integrin-collagen interactions.

**Fig. 7.**
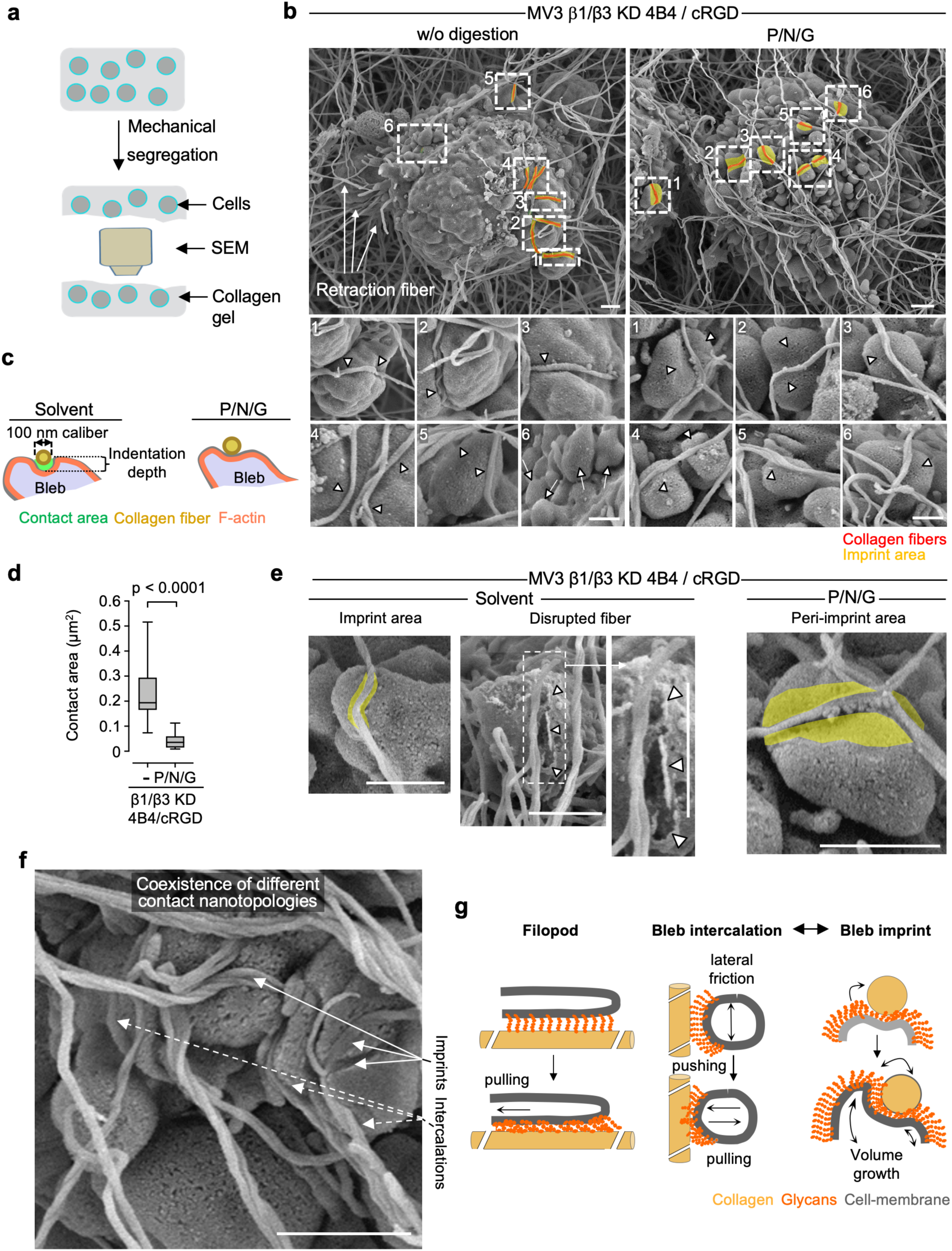
Morphology of glycan-dependent cell-matrix interactions. Scanning electron microscopy of MV3 β1/β3KD cells before and after glycan removal with P/N/G glycosidase cocktail. (a) Schematic representation of the sample preparation for SEM analysis. (b) Morphology of MV3 β1/β3KD cell surface interactions with collagen fibrils without (left panel; zoom 1-5 as representative interactions) or after treatment with P/N/G glycosidases (right panel; zoom 1-6) in 3D collagen lattices. Deformed contact region between cell membrane (colored yellow, arrowheads) and collagen fiber (colored red). (b, left panel; zoom 6) Representative position of contact-free blebs with fully spherical morphology for cells without P/N/G glycosidase treatment. Bar, 1 µm (b, overviews), 0.5 µm (b, ROIs). (c) Schematic representation of the imprint formation between a membrane bleb and a collagen fiber for cells without glycan removal and its loss after interference with cell surface glycosylation. (d) Area of membrane-collagen fiber contacts. Box and whisker plots show 25-75 percentiles (box), the median (middle line) and and 5/95 percentiles (whiskers). P value, non-paired Mann-Whitney test, 2 tailed. (e) ROIs showing collagen fibers which form deep imprints into membrane blebs of MV3 β1/β3KD cells (e, left panel, b, left panel; region 1-6) which are absent in cells after P/N/G glycosidase digestion (e, right panel, b, right panel; region 1-6). Bars, 0.5 µm. (f) Representative ROI showing coexistence of imprints and intercalations in MV3 β1/β3KD cells without glycan removal. Bar, 1 µm. (g) Glycans serve as multivalent and universal adhesion scaffold. Schematic representation of the glycocalyx-mediated adhesion strategies developed by cells after interference with β1/β3 integrin expression and function.

To test, whether the glycocalyx engages with substrate other than collagen BSA was used as non-specific ligand. Whereas integrins did not favor cell binding to BSA, glycans supported interaction of MV3 control, MV3 β1/β3KD and Molt-4 cells to BSA with forces similar to collagen binding (Fig. 4g, Extended Data Fig. 6h, 7b). This implicates the glycocalyx as an adhesion system to ECM and other substrates.

### Glycan-mediated interactions to collagen fibrils

We finally aimed to identify the cell contact structures mediated by surface glycans towards collagen fibers and first classified the protrusion types in control cells and cells after limiting integrin and glycan availability. Cells with polarized elongated or rounded morphology developed either (i) pointed actin-rich pseudopod- and filopod-like extensions or (ii) bleb-shaped, rounded protrusions in a cell-type dependent manner (Fig. 5a). After targeting integrins and converting from elongated to rounded morphology MV3 cells preferentially developed bleb-like protrusions (Fig. 5a, b, Supplementary Video 7), and this was similar to the bleb-rich morphologies of untreated migrating Molt-4 cells (Fig. 5b, Supplementary Video 8). By contrast, migrating β1-/-MEFs predominantly formed filopod- and bleb-like protrusions (Fig. 5a, b). Notably, both well-developed blebs (MV3, Molt-4) and filopod-like protrusions (MEFs) were diminished after additional glycan removal (Fig. 5a, b, Supplementary Video 7, 8), indicating that both protrusion types depend on an intact glycocalyx.

To identify the topologies and molecular organization of glycocalyx-mediated interactions to collagen fibers, we performed 3D confocal microscopy in fixed cells after integrin targeting and detected the distribution of surface glycans by extracellular applied lectins (ConA or MAA), together with the underlying actin cytoskeleton and the position of collagen fibers identified by reflectance. Filopod-like protrusions formed a linear glycan-rich interface along collagen fibers (Fig. 5c, d, left panels). Likewise, cell blebs were covered by a non-focalized glycan layer and formed complex surface topologies in contact with collagen fibers (Fig. 5c, d, right panels). Unexpectedly, at confocal microscopic resolution, the lectin signal surrounding bleb-like protrusions locally overlapped with cross-sectioned fibers in MV3 (Fig. 6a, upper and middle panel; solid arrows) and Molt-4 cells (Extended Data Fig. 8a, upper and middle panel; solid arrows). We applied quantitative image analysis of the position of collagen fibers relative to the outer rim of the glycocalyx and identified an overlap range of 50 to >200 nm in depth (Fig. 6b, c, Extended Data Fig. 8b), and this overlap was ablated after glycan removal (Fig. 6c, Extended Data Fig. 8b). As outcome, small blebs resided adjacent to collagen fibrils without colocalization (Fig. 6a, lower panel; Extended Data Fig. 8a, lower panel; dashed arrows). This indicates that the glycocalyx provides very tight apposition or even superimpose with the ECM structures.

To detect the 3D topologies of glycan-mediated contacts at high resolution, beyond the diffraction limit of light microscopy, we used scanning electron microscopy to visualize the surface of cells located inside the collagen matrix, by mechanically separating the matrix at mid-level (Fig. 7a). Blebs in direct contact with a collagen fiber formed complex-shaped indentations (Fig. 7b, left panel; arrowheads in regions 1-5), whereas contact-free blebs remained spherical and without indentation (Fig. 7b, left panel; arrows in region 6). The “grip-like” tight interphase between cell surface and collagen fibrils corresponded to a contact area ranging from 0.18 up to 0.29 µm^2^, calculated from the bleb diameter, fiber caliber and indentation depth, and the length of the interactions (Fig. 7c, d). Notably, these interactions withstood mechanical processing of the collagen lattice during sample preparation in most cells (Fig. 7e; yellow colored area). However occasional disruptions of the interaction uncovered a half-cylindrical grove matching the orientation and calibre of the displaced fibril (Fig. 7e, left panel; arrowheads), thus confirming the curved shape and otherwise tight submicron scaled apposition of the cell surface to the fibril (“nanogrips”). After enzymatic glycan removal, the nanogrips were perturbed in shape and lacked tight folding but instead showed a flattened, weakly concave area adjacent to the fiber (Fig. 7b, right panel; arrowheads in regions 1-6 and 7e, right panel; yellow colored area). This data indicates a previously unappreciated scaffold function of the glycocalyx, mediating grip-like membrane topologies towards irregular-shaped extracellular structures (Fig. 7f).

## Discussion

These results identify the glycocalyx in mediating low-to-moderately adhesive mechano-coupling in the pN force range and cell migration in 3D ECM, and neither of these functions depends on integrins. Glycan-mediated substrate interactions consist of linear membrane appositions to collagen fibrils of finger-like protrusions or curved membrane indentations across or adjacent to blebs (Fig. 7g). The graded transition between migration modes after interference with integrins and surface glycans indicates that integrin-mediated high- and glycan-mediated low-adhesive interactions coexist substitute for each other during migration and support migration plasticity^3^.

To detect the adhesion forces provided by surface glycans and simultaneously minimize overlapping integrin-mediated attachment, we antagonized integrin functions in adhesion and migration by combining downregulation, antibody and small molecule targeting, or genetic deletion. The obtained force spectra indicate that glycan- and integrin-mediated mechanocoupling in mesenchymal cells is additive and occurs in parallel, with ∼50% of adhesion to collagen mediated by β1 and β3 integrins, ∼30% by the glycocalyx, and residual attachment by yet unidentified bonds.

Integrin-mediated adhesions transmit strong mechanotransduction which underlie long-range ECM deformation and stiffening, and both are hallmarks of mesenchymal migration^32,33^. Strong adhesion in the nN range and spindle-shaped cell elongation were lost after inhibiting integrin functions but filopod- and bleb-like protrusions persisted and maintained polarization and movement in rounded cells, similar to leukocytes and mesenchymal cells moving within engineered non-adhesive environments without engaging integrins^9,29^. Thus, glycan-based adhesions towards collagen fibrils become apparent as a secondary actin-rich adhesion system when integrins are downregulated or disabled. Surface glycans provide a very tight interface between the cell membrane and hydrophilic surfaces, here collagen fibrils which is mechanically sufficient to mediate cell migration, and diverse physicochemical interaction may be at work, including electrostatic bonds, hydrogen bonds, Van der Waals forces^19,34^ and, possibly, Casimir forces^35^.

The resulting lattice-like bonds were associated with at least two distinct protrusion types and topologies including filopod-like linear-shaped appositions or curvature-based nanogrips. Both interaction types maintain a glycan-rich lattice between the cortical actin and collagen fibrils and both types were lost after enzymatic removal of the glycocalyx. In adhesive cells of mesenchymal differentiation, membrane folds of similar topology were shown to provide 3D shape alignment, intense integrin clustering to fibrillar collagen and strong force transmission^36^, whereas the here described glycocalyx-mediated membrane folds are developed by cells after integrin interference and lack focalized adhesion and cytoskeletal organization. Thus, both integrins and the surface glycocalyx can induce membrane folds to engage with collagen fibrils^36^. Integrin- and glycocalyx-dependent membrane folds likely represent complementary molecular principles to provide an adaptive range of adhesive membrane topologies that vary in shape, strength, lifetime and ECM contexts. Whether adhesions mediated by glycans also contribute to friction, by mediating bonds between flat surfaces and the cortical actin cytoskeleton and which maintains migration in other systems^9,10,23^, remains to be determined. Similarly, the link between glycan-mediated membrane apposition to collagen fibrils and other cell-surface dependent physical interactions with ECM discontinuities, such as elbowing, intercalation and propulsive shape change^11^, remain to be clarified. Since integrin- and glycan-mediated mechanotransduction likely occur in parallel, modulation of each adhesion system may adjust cell-matrix interaction and migration programs, including the mesenchymal-to-amoeboid transition.

## Materials and Methods

### Cells

Metastatic human MV3 melanoma cells stem from a spontaneous lung metastasis after subcutaneous implantation in nude mice^37^. For generation of β1/β3 integrin double knockdown cells shRNA sequences targeting ITGB1 (β1 integrin; AGCCACAGACATTTACATTAAA) and ITGB3 (β3 integrin; AAGTCACTTTCTTCTTCTTAAA) for gene silencing by RNA interference were cloned into the lentiviral vector pLBM either containing a puromycin (p-puro) and a neomycin (p-neo) cassette. Lentiviral particles were produced and concentrated by ultracentrifugation, as described^38^. MV3 parental cells were infected with p-puro and p-neo viruses (vector controls), or with ITGB1 (on p-puro) and additionally ITGB3-targeting (on p-neo) pLBM viruses. Stable MV3 β1/β3KD cells were maintained in medium supplemented with puromycin (5 µg/mL) and G418 sulfate (400 mg/mL). β1KD efficiency determined by Western blot using detection Ab EP1041Y (10 µg/mL) from whole cell lysates or flow cytometric analysis of β1 integrin surface levels. For interference with integrin-mediated adhesion for migration and force spectroscopy of collagen, anti-β1 integrin mAb 4B4 (10 µg/mL) and cRGD peptide (2 or 10 µM) in addition to stable β1/β3 integrin knockdown were used.

Stable Lifeact-eYFP expressing MV3 melanoma cells were obtained as follows. A fragment spanning the pMSCV-hygro (Addgene, 634401) Hygromycin B resistance cassette was amplified by PCR (primer sequences available upon request) and inserted into KpnI-digested pLenti6.2/V5-DEST (Invitrogen, V36820), replacing the Blasticidin resistance cassette (pLenti6.2_Hygro/V5-DEST). A BglII-NotI fragment, containing the Lifeact-eYFP open reading frame, was then isolated from plasmid pEYFP-N1-ΔATG-Lifeact^39^ and cloned into BamHI/NotI-digested pENTR4 (Addgene). Using Gateway recombinase-mediated transfer the pENTR4-Lifeact-eYFP insert was subsequently introduced into pLenti6.2_Hygro/V5-DEST. Stable Lifeact-eYFP expressing MV3 cells were obtained following lipofectamine 2000 transfection of the resulting plasmid, subsequent hygromycin B (Invitrogen) selection (200 μg/mL) and final fluorescence-activated cell sorting.

Immortalized floxed β1 (β1(fl/fl)), β1-/- and stably full-length β1/GFP fusion protein expressing murine embryonic fibroblasts (MEFs) were obtained as described^40^ and maintained in culture at 33 °C. Expression levels of β1 integrins in floxed β1 (β1(fl/fl)) cells and the loss of β1 expression in β1-/-cells were monitored by flow cytometry with mAb KMI6 (Extended Data Fig. 2a). For β1 integrin rescue experiments, MEFs stably expressing full-length β1/GFP fusion protein^41^ (Extended Data Fig. 2b) were analyzed.

The human T lymphoma cell line Molt-4 was kindly provided by Blanca Scheijen, Laboratory of Pediatric Oncology, Radboud Institute for Molecular Life Sciences, Nijmegen, The Netherlands. Unless stated otherwise, MV3 and Molt-4 cells were maintained in RMPI 1640 (Gibco, 21875-034) or DMEM (Gibco, 11965-092) (MEF) containing 10 % fetal bovine serum (FBS, PAA Laboratories, A15-101), penicillin/streptomycin 100 U/mL penicillin and 100 μg/mL streptomycin (Gibco, 15140-122), L-glutamine (4mM, PAN Biotech, P04-80100), sodium pyruvate (1 mM, Invitrogen, 11360) and detached from the culture plate by EDTA (2 mM, PAN Biotech, P10-026500) in 1X PBS (Gibco, 14040-117). Cell lines were tested routinely for mycoplasma contamination.

### Animals

Balb/c nu/nu mice (CAnN.Cg-Foxn1nu/Crl) were purchased from Charles River, Germany.

### Glycan removal

Adherent cells from subconfluent cultures (MV3, MEF) were detached using 2 mM PBS/EDTA. Molt-4 suspension cells were harvested directly. After washing, cells in RPMI1640 with 1% v/v FBS medium were incubated with glycosidase cocktail (P/N), containing hyaluronidase at 275-500 U/mL (Sigma Aldrich, H3506), heparitinase I at 10 mU/mL (Seikagaku, 100704), chondroitinase ABC at 100 mU/mL (Sigma Aldrich, C3667) and neuraminidase type V at 100 mU/mL (Sigma Aldrich, N2875) (37 °C, 5 % CO_2_, pH 7.0-7.4, 6 h). P/N targets distal glycosaminoglycan chains of proteoglycans (P) and terminal sialic acids / neuraminic acids (N) of glycoproteins and glycolipids. Alternatively, P/N was supplemented with β1,4 galactosidase at 150 mU/mL (QA-Bio, E-BG07), cleaving non-reduced terminal β1-4 galactose (G) of glycoproteins and glycolipids. Cells were used for functional studies without washing steps to prevent the loss of dead cells by centrifugation.

Because latent cytotoxicity may negatively impact cell adhesion and migration, cell viability was routinely monitored. After glycan removal cells were collected without centrifugation, stained by propidium iodide (2.5 µg/mL), and analyzed by flow cytometry. Viability was further derived from intact nuclear morphology after fixation in situ and DAPI staining. Strategies to delay glycan recovery, including inhibition of protein export by brefeldin A and prevention of protein glycosylation by tunicamycin, directly impacted migration efficiency and mode (data not shown), which precluded their use.

### Flow cytometry

Flow cytometry was performed to detect integrins and glycans at the cell surface. Cells were isolated from liquid culture using detachment by PBS/EDTA, washed in 1X PBS and transferred into 96 well plates, spinned down by centrifugation at 220 g, 5 min and washed three times in 1X PBS. Washing steps were excluded for cells used in glycosidase treatment experiments to prevent the loss of dead cells by centrifugation. For labelling of glycosaminoglycans (GAG), cells were incubated with anti-GAG single-chain antibodies from periplasmic fractions (HS4C3, 5 µg/mL in PBS; LKN1 and I03H10, 20 µg/mL in 1X PBS) for 45 min at 20 °C. Following three washing steps in PBS, 4 °C, cells were incubated with secondary antibody (P5D4, 10 µg/mL in PBS) from hybridoma culture supernatant for 30 min at 4 °C on ice, washed and incubated with the third antibody, goat-anti-mouse IgG Alexa Fluor 488 (10 µg/mL) for 30 min at 4 °C on ice. For lectin labelling, cells were incubated with MAA-biotin (100 µg/mL) in 1X PBS for 20 min at 4 °C on ice. Following the three washing steps in PBS, 4 °C, cells were incubated with Streptavidin-Alexa Fluor 488 (Invitrogen, S32354, 10 µg/mL) and incubated for 30 min at 4 °C on ice. Bioorthogonal metabolic labeling was used to detect cell surface sialic acids / neuraminic acids as described^42^. Cells were cultured for 6 days in the absence or presence of 50 µM peracetylated N-azidoacetylmannosamine (Ac4ManNAz). Medium and compound were changed after 3 days. Adherent cells were detached using 2 mM PBS/EDTA, washed in 1X PBS three times at 220 g for 5 min, resuspended in 1X PBS and incubated in 60 µM biotin-conjugated Bicyclo [6.1.0] nonyne (BCN-biotin) or buffer for 1h at 20 °C, washed three times, resuspended in ice-cold PBS containing Streptavidin-Alexa Fluor 488 (Invitrogen, S32354, 10 μg/mL) and incubated for 30 min at 4 °C. Cells were washed by centrifugation, resuspended in ice-cold PBS and transferred into conical FACS tubes (Micronic, M32,000). For harvesting cells from 3D collagen lattices, the collagen matrix was dissolved using 1000 U/mL collagenase I (Sigma Aldrich, C0130) for 30 min, 37 °C and cells were obtained without washing steps to prevent the loss of dead cells by centrifugation. Cell-associated fluorescence was measured by flow cytometry (FACSCalibur, BD Biosciences) and data were analyzed using FCS Express (Version 5 and 6 Research Edition; De Novo Software, Los Angeles, CA). Per sample, 10,000 morphologically intact and alive cells were gated (blue boxes) based on forward and sideward scatter (FSC/SSC plot) and propidium iodide negativity (FSC/Propidium iodide plot) was used to exclude unspecific fluorescence from the final signal intensities (Extended Data Fig. 1g). The same gating strategy was used to quantify the fraction of viable cells, after glycosidase treatment.

### Transmission electron microscopy (TEM)

Detection of the surface glycocalyx using TEM was performed largely as described^43^. MV3 cells were detached from the culture plate (EDTA), pelleted and fixed with 0.0075 % w/v Ruthenium Red (Sigma Aldrich, R2751) containing 3% v/v glutaraldehyde (Roth, UN-32658 III) for 1h at RT and 1% v/v osmium tetroxide (Roth, UN-24716.1 I) for 2h at RT in 0.1 M PHEM buffer (PIPES, HEPES, EGTA, MgCl_2_). Cells were washed with 0.1 M PHEM buffer, resuspended in 0.1 M PB and centrifuged (15,871g, 30 sec). The supernatant was discarded and pre-warmed 4 % w/v agar was applied to the cell pellet followed by an incubation (2-5 min, 45 °C). After centrifugation cell pellets were chopped up into slices and incubated in 2 % v/v paraformaldehyde / 0,1 M PB (2 h, RT). Samples were rinsed in 0.1 M PB, dehydrated through a graded ethanol series (50 %, 70 %, 80 %, and 96 % respectively for 5 min each and 2x 15 min in ethanol 100 % p.a.) and embedded in an epoxy resin. Samples were sectioned (50-80 nm), stained with 4% v/v uranyl acetate (20 min, RT), washed with Mill-Q, stained with lead citrate solution (2.66% w/v lead nitrate and 3.52 % v/v sodium citrate, 10 min, RT) again washed with Mill-Q, dried and examined by TEM (Jeol 1010, Jeol USA Inc., Peabody, USA). Statistical analysis was obtained using the non-paired t-test, 2 tailed.

### Scanning electron microscopy (SEM) of 3D cell-ECM contacts

MV3 cells in 3D bovine fibrillar collagen after 90 min culture (37 °C, 5% CO_2_) were fixed in 2% glutaraldehyde in 0.1M Cacodylate buffer for 1h at 37 °C, washed, post-fixed for 1h in 1 % OsO4 in 0.1M Cacodylate buffer, dehydrated by a graded ethanol series and critical point dried. Samples were mounted on stubs, upper part of collagen was mechanically pulled off with a tweezer, samples were coated with 5nm Chromium (Quorum Q150TS, Quorum Technologies Ltd, East Sussex, UK) and analyzed by SEM (Sigma 300, Carl Zeiss AG) operating at 3 kV.

The type and geometry of the interactions between cell surface and collagen fibers was quantified by digital image segmentation using Fiji. To determine the contact area, the lateral surface area of the fiber was calculated, and the bleb-free fiber area subtracted. Statistical analysis was obtained by the non-parametric, non-paired, Mann-Whitney t-test, 2 tailed.

### Atomic force Spectroscopy (AFS)

The forces between carbohydrate polymers and collagen fibers were determined by atomic force spectroscopy (AFS). Flat collagen lattices (1.6 mg/mL, 40 µm thick) mounted on a 1.5 mm thick glass cover slip (Plano, L43382) were probed using a NanoWizard AFM (JPK Instruments) mounted on an Axiovert 200 inverted microscope (Carl Zeiss AG).

The cantilevers were functionalized with either bovine serum albumin (BSA, 3%, 2 h, 20 °C, Sigma-Aldrich, A7906), Bis-NHS-polyethylen glycol (5 % w/w, 200,000 g/mol, Nektar), amylose (carboxymethyl amylose, Sigma Aldrich, C4947) or cellulose (carboxymethylcellulose, Sigma Aldrich, 419338) at the indicated weight percentage. After UV irradiation (10 min), the silicon nitride cantilevers were functionalized with (3-glycidyloxypropyl)trimethoxysilane (98%, Sigma Aldrich, 440167) at 80 °C for 30 min, incubated in sodium-borate buffer for one hour at RT, functionalized with NH2-PEG(3000)-COOH (50 mM, Rapp Polymers, 133000-20-32) in sodium borate buffer for 1h at RT. Then 1 % (w/w) BSA was coupled to the PEG (3000) for 1h at RT with 1-ethyl-3-(3-dimethylaminopropyl) carbodiimide (EDC, 100 mM, Sigma Aldrich, 39391) and N-hydroxy-succinimide (NHS, 100 mM, Sigma Aldrich, 130672). The measurement was performed without BSA in solution. After UV irradiation (10 min), silicon nitride cantilevers (MLCT, Veeco Probes) with a spring constant of 60 pN/nm were functionalized by N1-(3-trimethoxysilylpropyl)diethylenetriamine (30 min, RT, Sigma Aldrich, 06666) followed by heating (30 min, 80 °C), incubation in sodium borate buffer (60 min; RT), and either Bis-NHS-PEG solution, amylose, or cellulose solution in N-hydroxysuccinimide (NHS; Aldrich; 100 mM) and 1-ethyl-3-(3-dimethylaminopropyl)carbodiimide (EDC, Sigma; 100 mM, 90 min, RT)^44^. System position and sensitivity were calibrated before each measurement, as described^45^.

The spring constants ranged from 50 to 80 (median 60) pN/nm (thermal noise method). All interaction measurements were performed in PBS/BSA (0.4%) at RT (set point of 200 pN for approach curve; dwell time 0.1 s). For each measurement, 1,600 to 10,000 force curves were obtained for a retract velocity of 10 µm/s from different spots within a 40 × 40 µm collagen surface area. Statistical analysis was obtained by the non-paired Mann-Whitney Wilcoxon test, 2 tailed using software package R, version 1.22 (R Foundation for Statistical Computing).

### AFM based single cell force spectroscopy (SCFS)

The interaction strength between cell surface glycans and 3D collagen I fibers was determined by AFM based single-cell force spectroscopy (AFM-SCFS), using a combined Catalyst AFM (Bruker) and inverted 3-channel Leica TCS SP5 II confocal laser-scanning microscope equipped with 10× 0.4 NA, 20× 0.70 NA, and 40× 0.85 NA air objectives and a brightfield camera (ORCA-05G, Hamamatsu).

Bovine collagen I (1.6 mg/mL) and BSA (2% in TSM solution, containing 20 mM Tris-HCl, 150 mM NaCl, 1 mM CaCl_2_, 2 mM MgCl_2_ at pH 8.0; incubated 24h at 4 °C) were used as substrates and prepared in Willco dishes (35 mm, Willco Wells B.V., HBSt-3522) as described^46^. Tip-less AFM cantilever (type D, Bruker, NP-O) with a nominal spring constant of 0.06 Nm^-1^ were coated with ConA (Sigma, C7275)52 and applied for single cell adhesion as published^47^. Cells were prepared as described above and maintained in medium containing 10 mM HEPES (Thermo Fisher Scientific, 15630080) to keep the pH constant between 7.4 and 7.5. Cell measurements were performed at 37 °C. Adhesion of the cantilever-bound cell to different substrates was measured after pushing the cell towards the substrate, applying 2 nN force for contact times varying between 0.5 to 60 s. Subsequently, the cell was retracted at 5 µm/s and allowed to recover for a time period equal to the precedent contact time^47^.

The maximum detachment force F [max] was calculated from the peak of the retraction curves to background level. Mean values were compared using a non-paired t-test, 2 tailed. The sequence of contact time measurements was varied and performed in different areas of the substrate. To rule out intermittent parameters, such as receptor activation, cytoskeletal reinforcement or epitope saturation due to receptor shedding, repeated probing conditions on one spot were verified and limited to 5 times in a row. AFS analysis was performed described^46^ using IGOR Pro 6.2 (Wavemetrics, Portland, USA) and Matlab R2012a and R2018 (MathWorks, Massachusetts, USA).

### Intravital multiphoton microscopy and image analysis

All intravital imaging experiments were approved by the Ethical Committee on Animal Experiments and performed in the Central Animal Laboratory of the Radboud University, Nijmegen (RU-DEC 2009-174, 2011-298), in accordance with the Dutch Animal Experimentation Act and the European FELASA protocol (www.felasa.eu/guidelines.php). Lifeact-eYFP expressing MV3 control and β1/β3KD cells were incubated with glycosidases for 6 h, at 37 °C followed by an incubation with adhesion perturbing anti-β1 integrin mAb 4B4 (10 µg/mL) and anti-αV integrin mAb 272-17E6 (10 µg/mL). Cells as single cell suspension (2×104 cells in 10 µL) were injected into the deep dermis of Balb/c nu/nu mice (Charles River) carrying a dorsal skin-fold chamber. To ensure integrin blocking from interstitial fluids, mice received i.p. injections of 4B4 (5 µg/g) and 17E6 (5 µg/g) 2 h before cell injection.

Intravital microscopy was performed 2 h after cell injection on anesthetized mice (1-3 % isoflurane in oxygen) on a temperature-controlled stage (37 °C). Blood vessels were labeled by intravenous injection of AlexaFluor750-labeled 70kD dextran (2 mg/mouse; Invitrogen). Imaging was performed on a customized near-infrared/infrared multiphoton microscope (TriMScope-II, LaVision BioTec, Bielefeld, Germany), equipped with three tunable Ti:Sa (Coherent Ultra II Titanium:Sapphire) lasers and an Optical Parametric Oscillator (OPO). 4D time-lapse recordings were acquired by sequential scanning with 960 nm (YFP, 10-20 mW) and 1090 nm (Al750 and SHG, 30-60 mW) with a sampling rate of 1 frame / 10 min over periods of up to 8 h. Images were processed using Fiji/ImageJ (http://pacific.mpi-cbg.de/wiki/index.php/Fiji, ImageJ, U. S. National Institutes of Health, Bethesda, USA). Drifts in time-lapse recordings were corrected using the StackReg plugin^48^. To avoid tissue regions perturbed by the injection procedure, migration analysis was performed in intact collagen-rich loose connective tissue identified by second harmonic generation. Quantitative analysis of tumor cell migration was performed using the FIJI manual tracking plugin (Fabrice P. Cordelières, rsb.info.nih.gov/ij/plugins/track/track.html). To control for small local tissue drifts, particularly important for analyzing the residual migration after glycan removal, movements of 3 tissue structures in close proximity to each analyzed cell were recorded and used to correct individual cell migration tracks.

### Statistical analysis

Statistical analyses were performed with GraphPad Prism 8 (GraphPad Software, USA), for AFS analysis additionally with software package R, version 1.22 (R Foundation for Statistical Computing), IGOR Pro 6.2 (Wavemetrics, Portland, USA) and Matlab R2012a and R2018 (MathWorks, Massachusetts, USA). Flow cytometry analyses were performed using FCS Express 5 and 6 (De Novo Software, Pasadena, USA). Computer-assisted cell tracking was performed using Autozell 1.0 software (Center for Computing and Communication Technologies [TZI], Bremen, Germany), Fiji/ImageJ (http://pacific.mpi-cbg.de/wiki/index.php/Fiji, ImageJ, U. S. National Institutes of Health, Bethesda, USA) and the FIJI manual tracking plugin (Fabrice P. Cordelières, rsb.info.nih.gov/ij/plugins/track/track.html). T-test and ANOVA were performed after data were confirmed to fulfil the criteria using the Shapiro-Wilk normality test, otherwise Kruskal-Wallis tests or Mann-Whitney U-tests were applied, and post-hoc correction (Bonferroni, Dunn) was performed when multiple samples were compared. The sample numbers and applied statistical analyses for all experiments are shown in Extended Data Table S1.

### Data availability

All data are available from the authors on request.

## Code availability

The used custom algorithms are available from the authors on request.

## Acknowledgements

We thank Monika Kuhn, Ute Eifert, Martina Jossberger and Margit Ott for expert technical assistance; Irmgard Schwarte-Waldhoff, Christian Linden, Kimberly Burns, and Ute Schwab for technical advice; Jan Schepens and Wiljan Hendriks for generating the Lifeact-eYFP expressing cells; Zena Werb for helpful discussions and Martin Lohse and Rex Chisholm for comments on the early version of the manuscript. The authors furthermore thank the Radboud Consortium for Glycoscience for helpful discussions and project support and Radboudumc Microscopic Imaging Center for the use of their facilities.

This work was supported by the Deutsche Forschungsgemeinschaft (DFG) (FR1155/8-1, 8-2 and 8-3 SPP-1190); an infrastructure grant to the Rudolf Virchow Center (FZ82); NWO-Vici (918.11.626), the European Research Council (617430-DEEPINSIGHT), the NIH (U54 CA210184-01) and the Cancer Genomics Center (CGC.nl) to PF; Kavli Institute at Cornell Postdoctoral fellowship (S.S.); the Wellcome Trust (grant 092015) and Cancer Research UK (grant C13329/A21671) (to M.J.H.), and the Center for NanoScience (L.R. and K.E.G) and the Fonds der Chemischen Industrie (Liebig Fellowship to K.E.G.). J.t.R. was supported by the Netherlands Organisation for Scientific Research (NWO) (Veni Grant No. 680-47-421). The AFM infrastructure was supported by NWO Medium Sized Investment (Grant No. NWO-ZonMW 91110007).

## Author contributions

S.S., M.J.H., D.J.L., K.E.G. and P.F. designed the experiments, S.S, P.F., V.t.B. and M.t.L. performed migration, immunostaining experiments, S.S. and P.F. performed flow cytometry, S.S and B.W, performed *in vivo* imaging experiments, S.S., L.R., K.E.G. and J.t.R. performed AFM measurements, S.S. and M.K.-V. performed AFM confocal microscopy, S.S. and P.F., M.t.L. and V.t.B. performed confocal microscopy, M.t.L., M.W.-R., S.S. and J.F. performed electron microscopy, B.L., S.S. and L.B. performed Surface Plasmon Resonance spectroscopy, A.J.M., U.M., and R.M. generated β1 integrin deficient MEFs, S.K. generated β1/β3 integrin KD MV3 cells; S.S., P.F., V.t.B. and M.t.L. analyzed the data; S.S. and P.F. wrote the manuscript; all authors corrected the manuscript.

## Additional information

**Extended Data Information** is available in the online version of the paper.

**Correspondence and requests for materials** should be addressed to P.F.

