## Supplementary Material for "Glycocalyx-mediated Cell Adhesion and Migration"

[1] Departments of Cell Biology and [2] Biochemistry of Integrated Systems, Radboud Institute for Molecular Life Sciences, [3] Department of Radiology and Nuclear Medicine, [4] Department of Tumor Immunology, [5] Department of Neurology, Translational Metabolic Laboratory, Donders Institute for Brain, Cognition and Behavior, Radboud University Medical Center, P.O. Box 9101, 6500 HB Nijmegen, The Netherlands; [6] Kavli Institute at Cornell for Nanoscale Science and [7] Robert Frederick Smith School of Chemical and Biomolecular Engineering, Cornell University, Ithaca, New York, USA; [8] Rudolf Virchow Center for Experimental Biomedicine and Department of Dermatology, University of Würzburg, Josef-Schneider-Str. 2, 97080 Würzburg, Germany; [9] David H. Koch Center for Applied Research of Genitourinary Cancers, University of Texas MD Anderson Cancer Center, Houston, Texas, USA; [10] Wellcome Trust Centre for Cell-Matrix Research, Faculty of Life Biology, Medicine & Health, Univ. of Manchester, Manchester M13 9PT, UK; [11] Institute for Applied Physics, Ludwig Maximilians University, 80799 München, Germany; [12] Institute for Experimental Physics, University of Ulm, 89069 Ulm, Germany; [13] Department of Medical Biotechnologies, University of Siena, 53100 Siena, Italy; [14] Laboratorio di Patologia Clinica, Azienda Ospedaliera Universitaria Senese, 53100 Siena, Italy; [15] Joslin Diabetes Center, Harvard Medical School, 02215 Boston, MA, USA: [16] Cancer Genomics Centre, 3584CG Utrecht, The Netherlands; [17] Department of Preclinical Imaging and Radiopharmacy, Eberhard Karls University Tübingen, Germany; [18] CRUK Cambridge Centre, University of Cambridge, Cambridge, CB2 0RE, UK

**Supplementary Materials and Methods**

**Reagents.** The following primary antibodies were used: FITC-conjugated mouse anti-human CD29 IgG1 mAb (4B4-FITC Beckman Coulter, 6603109), mouse anti-human CD29 IgG1 mAb (4B4 Beckman Coulter, 6603113); FITC-conjugated mouse anti-human CD29 IgG2a mAb (K20, Beckman Coulter, IM0791U); rabbit anti-human CD29 IgG mAb (EP1041Y, Millipore, 041109); mouse anti-human CD18 IgG1 mAb (7E4, Beckman Coulter, IM1567); FITC-conjugated mouse anti-human CD18 IgG1 mAb (7E4, Beckman Coulter, IM1568); mouse anti-human CD51 IgG1 mAb (272-17E6, Abcam, ab16821); FITC-conjugated mouse anti-human CD51 IgG1 mAb (AMF7, Beckman Coulter, IM1855); mouse anti-human CD61 IgG1 mAb (Y2/51, Bio-Rad AbD Serotec, MCA2588GA); FITC-conjugated mouse anti-human CD61 IgG1 mAb (SZ21, Beckman Coulter, IM1758); rat anti-human CD104 IgG2 mAb (439-9B, BD Biosciences, 555719); FITC-conjugated rat anti-human CD104 IgG2 mAb (439-9B, BD Biosciences); rat anti-human β7 integrin IgG2 mAb (FIB504, ThermoFisher Scientific, 14-5867-82); PE-conjugated rat anti-human β7 integrin IgG2 mAb (FIB504, BD Biosciences, 555945); mouse anti-human CD49c IgG1κ mAb (C3II.1, BD Biosciences, 556024); mouse anti-human CD49d IgG1 mAb (9F10, BD Biosciences, 555502); rat anti-human CD49f IgG2 mAb (GoH3, BD Biosciences, 555734); rat anti-mouse CD29 IgG mAb (KMI6, BD-Pharmingen, 558741); rat anti-mouse CD18 IgG2a mAb (M18/2, BD Biosciences, 557437); Armenian hamster anti-mouse CD51 IgG mAb (HMαV-1, Biolegend, 153202); rabbit anti-mouse CD61 IgG polyclonal Ab (Cell Signalling Technology, 4702); rat anti-mouse CD104 IgG2a mAb (346-11A, Biolegend, 123602); rat anti-mouse β7 integrin IgG2 mAb (FIB504, BD Biosciences, 555943); rabbit anti-human GAPDH polyclonal Ab (Sigma-Aldrich, G9545). Phage-display anti-GAG single-chain antibodies from periplasmic fractions carrying a vesicular stomatitis virus glycoprotein (VSV-G)-tag for recognition by a secondary antibody (kind gift from Toin van Kuppevelt, Radboudumc, Nijmegen), including anti-heparan sulphate (HS) antibody (HS4C3)[49,50], anti-dermatan sulphate (DS) antibody (LKN1)[51] and the anti-chondroitin sulphate (CS) antibody (I03H10)[52]; rabbit anti-human discoidin domain receptor (DDR1) IgG pAb (C-20, Santa Cruz BioTech, sc-532); rabbit anti-human discoidin domain receptor (DDR2) IgG pAb (H-108, Santa Cruz BioTech, sc-8989); mouse anti-human CD138 IgG1 mAb (B-B4, Bio-Rad AbD Serotec, MCA681H); FITC-conjugated rat anti-CD44 IgG2bκ mAb (IM7, Thermo

Fisher Scientific, 11-0441-85); rat anti-human CD44 IgG2a mAb (Hermes-1, Endogen, MA4400)[53]. The following isotype controls were used: Mouse IgG1 mAb-1 (NCG01, Neomarkers, NC-748-PABX); mouse IgG1κ mAb (MOPC-21, BD Biosciences, 555746); rat IgG2bκ (R35-38, BD Biosciences, 555846); FITC-conjugated rat IgG2bκ (A95-1, BD Biosciences, 556923); FITC-conjugated rat IgG2bκ mAb (eB149/10H5, Thermo Fisher Scientific, 11-0441-85); rat IgG2aκ mAb (R35-95, BD Bioscience, 555841); FITC-conjugated rat IgG2aκ (R35-95, BD Bioscience, 553929); rabbit IgG pAb (R&D Systems, AB-105-C); Armenian hamster IgG (HTK888, Biolegend, 400901). The following secondary and tertiary antibodies were used: Mouse anti-vsv-g (clone P5D4) IgG1 mAb from hybridoma culture supernatant recognizing the vsv-g tag (kind gift from Toin van Kuppevelt, Radboudumc, Nijmegen); IRDye 800CW-conjugated goat anti-rabbit IgG Secondary Antibody (Li-Cor, 926-32211); Alexa Fluor 488-conjugated goat-anti-mouse IgG (H+L) (Thermo Fisher Scientific, A-11029); FITC-conjugated goat-anti-mouse IgG (H+L) (Thermo Fisher Scientific, 62-6511); Alexa Fluor 488-conjugated goat-anti-hamster IgG (H+L) (Thermo Fisher Scientific, A-21110); Alexa Fluor 488 conjugated goat anti-rat IgG (H+L) (Invitrogen, A-11006); Alexa Fluor 488-conjugated goat-anti-rabbit IgG (H+L) (Invitrogen, A-11008). The following lectins and secondary fluorophores were used: Tetramethylrhodamine-conjugated concanavalin A (ConA-TMR, Invitrogen, C860), biotin-labeled Maackia amurensis agglutinin (MAA-biotin, EY Laboratories, BA-7801-5), Alexa Fluor 488 or 647-conjugated streptavidin (Invitrogen, S32354, S32357). Alexa Fluor 488 conjugated phalloidin (Invitrogen, A12379), cyclic tripeptide arginine-glycine-aspartic acid (cRGD; Arg-Gly-Asp-D-Phe-Val) and cRAD control peptide (Arg-Ala-Asp-D-Phe-Val) (purity >95 %, Bachem; H-2574, H-4088), 4',6-diamidino-2-phenylindole (DAPI, Sigma Aldrich, D9542) was used for staining of nuclei. Propidium iodide (Sigma Aldrich, P4170) was used for monitoring cell viability.

**Protein gel electrophoresis and Western blot.** Protein β1 integrin content and efficient knockdown was determined by protein gel electrophoresis and Western blot analysis using whole cell lysate and incubation with β1 integrin antibody (2 μg/mL) and GAPDH antibody (1:10,000) as loading control and fluorescence detection using and Odyssey CLX imaging system (Li-Cor) followed densitometric analysis of the relative signal reduction.

**Cell sorting.** Cell sorting was performed using the FACS Elite cell sorter, Beckman Coulter, Pasadena, USA). Sorted cells were used for migration and confocal analysis. MV3 cells were detached by PBS/EDTA, washed with 1X PBS, stained with anti-β1 integrin mAb 4B4 (10 µg/mL) and secondary AlexaFluor 488-conjugated goat-anti-mouse IgG antibody (10 µg/mL; see point 5. above), resuspended in cold culture medium, transferred into conical screw cap Falcon tubes (Thermo Fisher Scientific, 352070) and subjected to sorting of cells without residual integrin expression. Post sort purity of MV3 β1 integrin KD cells was assessed by flow cytometry, and a second round of cell sorting was used to eliminate occasional integrin-high cells. Accordingly, only morphologically intact cells with uniformly low residual integrin expression cells were included in experiments.

To prevent an impact on actin dynamics and cell migration, cell subsets expressing high Lifeact-eYFP levels were excluded by cell sorting. Lifeact-eYFP transfected MV3 melanoma cells were detached by PBS/EDTA and washed in PBS, resuspended in pre-warmed culture medium, transferred into conical screw cap Falcon tubes (Thermo Fisher Scientific, 352070) and subjected to sorting of eYFP positive cells with intermediate expression levels. Purity and uniformity of MV3 Lifeact-eYFP expressing cells post sorting were assessed by flow cytometry. The quality of the sorting was further verified by examining fluorescence intensity in live cells by confocal microscopy.

**Cytotoxicity tests.** Cell viability was determined by the CellTiter-Glo® ATP luminescent cell viability assay (Promega, G7570) as specified by the supplier. Briefly, MV3 β1/β3KD and Molt-4 cells were incubated with glycosidase cocktail P/N/G at 37 °C, 5 % $CO_2$ for 6 h. Following digestion 75,000 cells in 100 µL were seeded per well in 96 well plates. Subsequently CellTiter-Glo® reagent was added in an equal volume of 100 µL per well. The well plates were placed on an orbital shaker and mixed for 2 minutes at room temperature to induce cell lysis. Cells were incubated for another 10 minutes at room temperature to stabilize the luminescent signal. Bioluminescence was measured using a multi-detection microplate reader (Synergy 2, BioTek Instruments) and analyzed using Gen5™ (Version 1.07, BioTek Instruments). Residual cells from the enzymatic digestion procedure were embedded into 3D pepsin-digested bovine dermis collagen lattices and incubated at 37 °C, 5 %

$CO_2$. For harvesting cells from 3D collagen lattices after 6 h and 24 h (MV3 β1/β3KD) and 5 h and 15 h (Molt-4), the collagen matrix was dissolved using 100 U/mL collagenase I (Sigma Aldrich, C0130) for 15 min, 37 °C. Cells were counted, adjusted to 150,000 cells per 100 µL and viability analysed using the CellTiter-Glo® assay.

**Cell migration in 3D collagen lattices, cell tracking and persistence measurements.** Integration of cells into 3D pepsin-digested bovine dermis collagen lattices containing 97% type I and 3% type III (PureCol; Advanced Biomatrix), time-lapse microscopy and computer-assisted cell tracking[14] and morphometric analysis of mesenchymal and amoeboid phenotypes[54] were performed as described. The collagen was native, based on its sensitivity to cleavage by MMP-14 and resistance to degradation by trypsin[55]. Cell migration experiments were performed at 37 °C (MV3, Molt-4, MEF parental and MEF β1-/- cells) for up to 108 h. Multicellular spheroids of MEFs were generated by overnight culture on dishes coated with poly(2-hydroxyethyl methacrylate) (0.33%; Sigma Aldrich, P3932), individually transferred by the tip of a pipette and incorporated into collagen solution prior to polymerization. The population speed was obtained by tracking of randomly selected cells for 24 h observation periods using computer-assisted cell tracking was performed using Autozell 1.0 software (Center for Computing and Communication Technologies [TZI], Bremen, Germany). Persistence was calculated as mean for each cell from the continuous-time quotient of direct distance between the endpoints divided by the actual track length over the indicated time period, excluding phases of immobilization.

**Confocal microscopy and image analysis.** Confocal fluorescence and reflection microscopy were performed as described[14,54]. Cells in 3D collagen lattices were fixed in de-polymerized paraformaldehyde (4% v/v, 37 °C, Sigma Aldrich, P-6148), washed, and stained with the appropriate fluorescent reagents. Confocal z-reconstruction was performed on a TCS SP8 confocal microscope (Leica Microsystems, Mannheim, Germany) equipped with an HCX PL APO 63x 1.2 NA water immersion lens or on a Leica SP2 scanner using a 60x 1.2 NA oil immersion lens. To minimize crosstalk in multi-fluorescence samples, each channel was acquired separately and displayed by maximum intensity projection of selected slices. Image analysis was performed using the Fiji ImageJ software from native slices (V1.51d V1.67 and 2.0, U. S. National Institutes of Health, Bethesda, USA).

To quantify the position of collagen fibers relative to the cell surface, confocal microscopy cross sections were used for analyzing the overlap (i.e. imprint depth) of the collagen fibers into the glycocalyx (MAA) or cortical actin layers (phalloidin) (Figure 6B). The position of the cell membrane directly flanking the cross sectioned fiber was cross-referenced with the position of the fiber relative to the cell surface by plotting their intensity profiles along measuring lines with a constant length of 8 µm and a width of 5 pixels at different positions. The obtained intensity profiles were averaged and plotted to derive the half-maximum intensities ($Max_{1/2}$) of MAA, phalloidin and collagen as reference points. The imprint depth (ID) was calculated based on the equation:

$$ID(nm) = \frac{(x\text{-value MFI}_{[Max1/2]} - x\text{-value Reflection}_{[Max1/2]})}{PSF}$$

with MFI representing the MAA or phalloidin signal and reflection the position of the collagen fiber. To correct for z-aberrations and assume point-like geometry, the positional values were divided by the point spread function in z-direction (PSF: 2.75).

**Surface plasmon resonance (SPR).** The binding affinities between amylose polymers and collagen fibers were monitored by surface plasmon resonance (SPR) using a Biacore T100 instrument (GE Healthcare). Unless stated otherwise, all materials were purchased from GE Healthcare. For the preparation of the analyte, amylose type III (Sigmal Aldrich) was dissolved using 200 mM NaOH and subsequently neutralized with 200 mM HCl. The resulting amylose solution was diluted in Biacore running buffer (HBSN, 10 mM HEPES, 150 mM NaCl, pH 7.4). For the preparation of the ligand collagen was immobilized via amino groups on the dextran surface of a CM5 sensor chip. The amylose solution at concentrations ranging from 300 µg/mL to 12 µg/mL was injected over the immobilized collagen or a control flow cell at the flow rate of 50 µL/min at 25 °C. Regeneration was achieved with a short pulse of 100 mM NaOH. No binding was observed with PEG dissolved in HBSN. The negligible non-specific binding on the empty flow cell was subtracted from measured specific binding affinity. Kinetics were analyzed using Langmuir model 1:1 for fitting (Biacore T100 evaluation 1.1.1 software). For the global fitting, a theoretical molecular weight of 8000 was chosen for amylose. In addition, the dissociation rates were calculated which is time-dependent and not influenced by the molecular weight of the analyte.

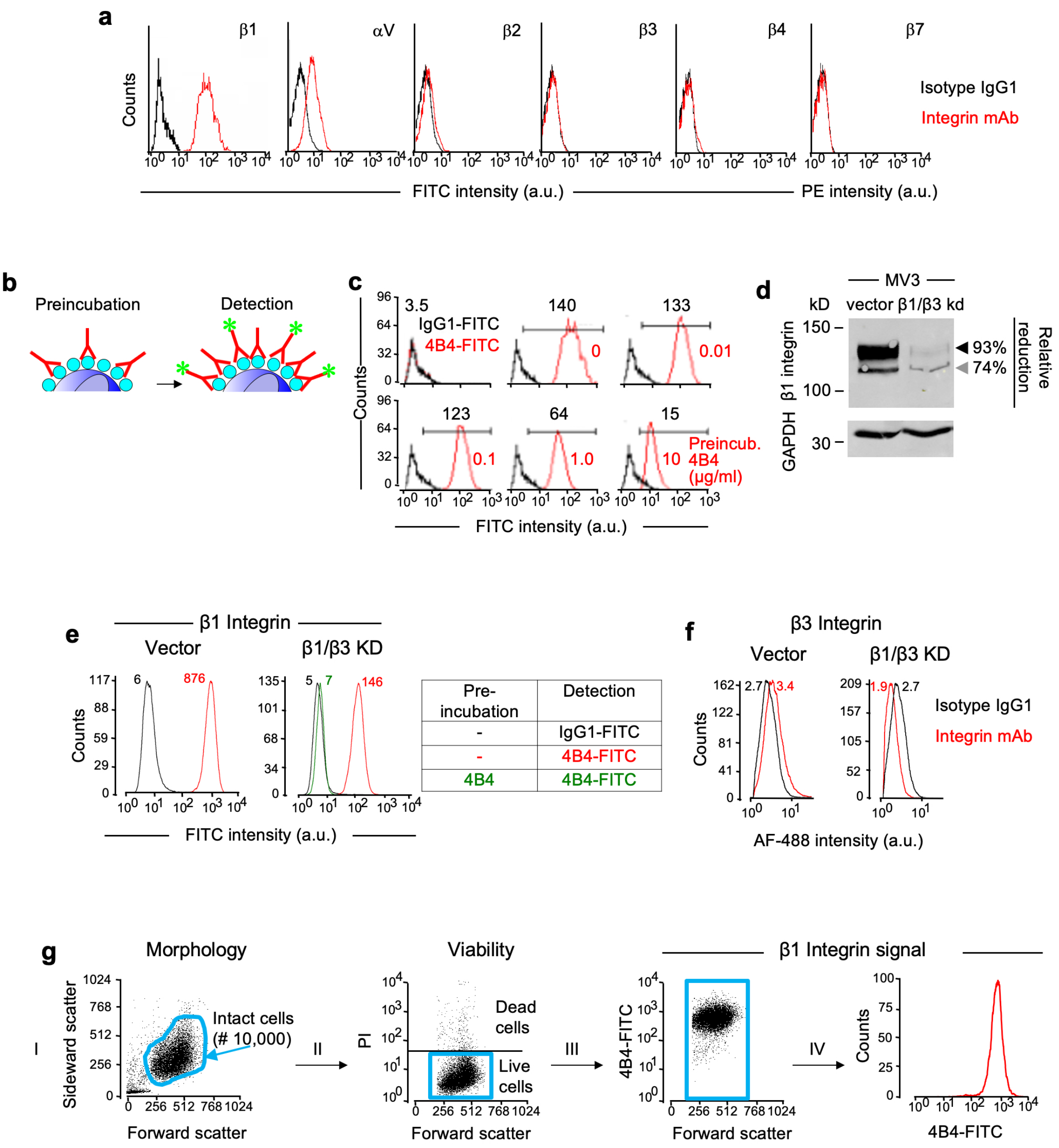


**Extended Data Fig. 1. Modulation of integrin expression and function in MV3 cells.** (a) Integrin surface levels on wild-type MV3 cells after detachment from subconfluent cultures. Red line, anti-integrin mAb; black line, isotypic control antibody. Data represent 1 experiment. (b-d) Epitope detection by fluorescent anti-β1 integrin mAb 4B4 in response to preincubation with unconjugated mAb 4B4. (b) Principle of the detection assay. MV3 cells were incubated for 30 min at 4 °C with non-fluorescent mAb 4B4 at different concentration, washed, stained with FITC-conjugated detection mAb 4B4 (10 µg/mL), and analyzed by flow cytometry. (c) Mean fluorescence intensity profiles of FITC-conjugated β1 integrin detection mAb 4B4 after pretreatment with non-labeled mAb 4B4 (red curves) or isotypic control antibody (black curve). Red numbers refer to the used 4B4 concentrations for pretreatment. (d) Knockdown β1 integrin detected by Western blot and (e) flow cytometry, reaching respective efficiencies of >90% and >80%. Full-length β1 integrin of vector control cells ran at approximately 135 kD (black arrowhead) and non-glycosylated pre-β1 at 90 kD[56] (gray arrowhead). GAPDH, 36-38 kDa. Unmodified scan of the Western blot is provided in Source data. (e) Saturation of residual epitope in MV3 β1/β3KD cells by additional incubation with anti-β1 integrin mAb 4B4, reaching approx. 97% reduction of available β1 integrin epitope detected by flow cytometry. (f) β3 integrin surface levels on MV3 vector control and β1/β3KD cells. Red line, anti-integrin β3 mAb Y2/51 (10 µg/mL); black line, isotypic control antibody (10 µg/mL). (E, F) Data represent 1 experiment. (g) Principle of the gating strategy used for the analysis of flow cytometry datasets. Per sample, 10,000 morphologically intact (I) and alive cells (II) were gated (blue boxes) based on forward and sideward scatter and propidium iodide negativity to exclude unspecific fluorescence from the final signal intensities (III, IV). Data represent 1 experiment. See also **Figure 1**.

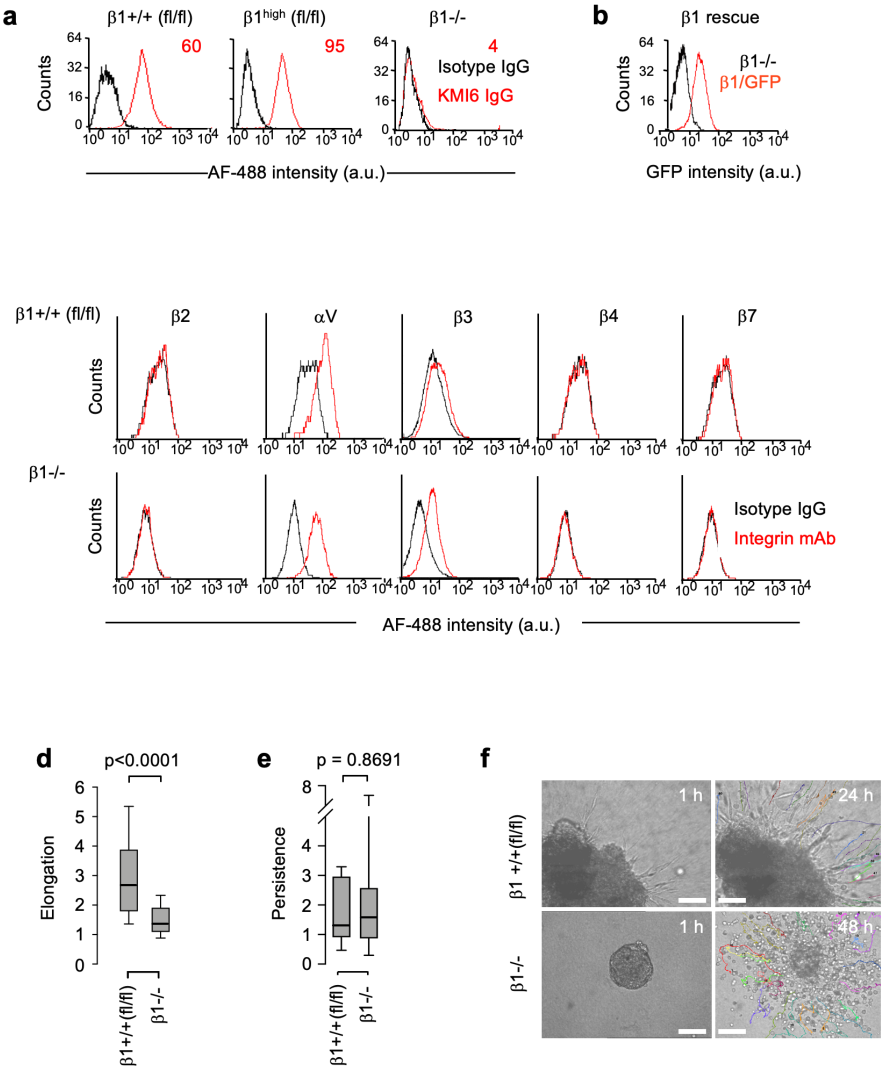


**Extended Data Fig. 2. Modulation of integrin expression and function in MEF cells.** (a) Flow cytometry analysis of β1+/+ (fl/fl), β1$^{high}$ (fl/fl) and β1-/- (fl/fl) MEFs. β1-/- MEFs lacked detectable β1 integrin, while the floxed β1 integrin expressing cells showed a distinct fluorescence signal. (b) Re-expression of β1/GFP in β1-/- MEFs, resulting in moderate expression level. (c) Surface expression of integrins on β1(fl/fl) and β1-/- MEFs. Except minor upregulation of β3 integrins, no further expression regulation in β1-/- MEFs was detected. (a-c) Data represent 1 experiment. (d-f) Mesenchymal-to-amoeboid transition and persistent migration in murine embryonic fibroblasts after genetic deletion of β1 integrin. (d) Elongation of β1(fl/fl) and β1-/- MEFs cells. Box and whisker plots show 25-75 percentiles (box), the median (middle line) and 5/95 percentiles (whiskers). P value, non-paired Mann-Whitney test, 2 tailed. Data represent 80 cells each from 2 independent experiments. (e) Persistence of β1-expressing and β1-/- MEFs. Data represent 11 β1+/+ (fl/fl) cells and 37 β1-/- cells from 3 independent experiments. Box and whisker plots show 25-75 percentiles (box), the median (middle line) and 5/95 percentiles (whiskers). P value, non-paired Mann-Whitney test, 2 tailed. (f) Migration of β1+/+(fl/fl) and β1-/- MEFs from multicellular spheroids into 3D collagen matrix visualized by bright-field microscopy (left) and individual cell tracking and display of migration paths (right). 1 representative out of 3 independent experiments. Bars, 10 μm. See also **Figure 1**.

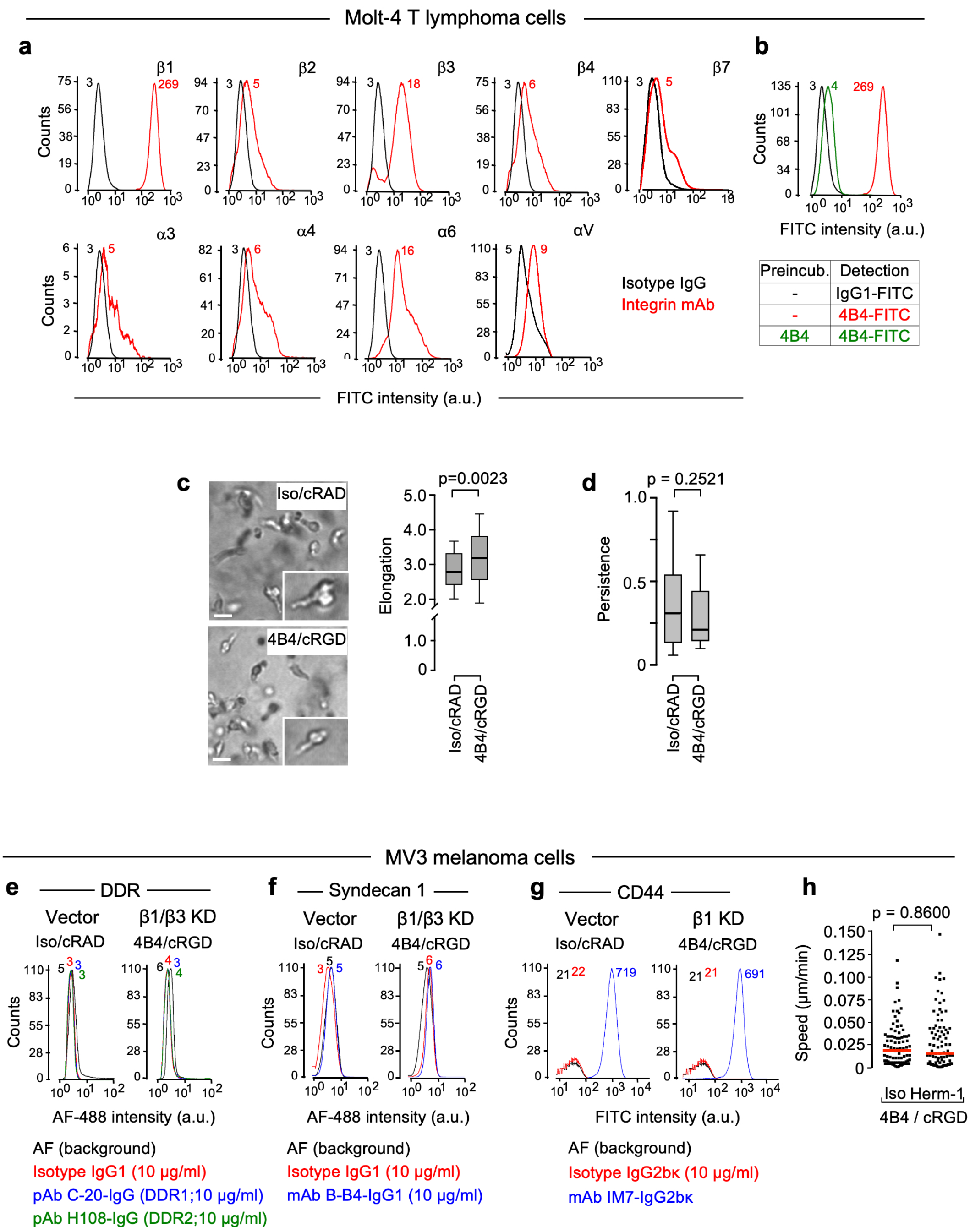


**Extended Data Fig. 3. Expression profile of integrins, alternative collagen receptors and migration phenotype of cells in 3D collagen matrix.** (a-d) Integrin expression profile and amoeboid phenotype and migration after interference with integrins in T lymphoma cells. (a) Integrin surface levels on Molt-4 cells. (b) β1 integrin epitope availability on Molt-4 cells after targeting of β1 with mAb 4B4 (10 µg/mL) detected by flow cytometry. Reduction of the 4B4 epitope was 97% (green line). Red line, anti-integrin mAb; black line, isotypic control antibody. (a, b) Data represent 1 experiment. (c) Morphology, elongation and (d) persistence of Molt-4 T lymphoma cells in the presence of β1 blocking antibody (4B4, 10 µg/mL) and β3 integrin masking cRGD (10 µM), compared with isotype and cRAD incubated control cells. Bars, 20 µm. (c) Box and whisker plots show 25-75 percentiles (box), the median (middle line) and 10/90 percentiles (whiskers). P value, non-paired Mann-Whitney test, 2 tailed. Data represent 99 cells each from 3 independent experiments. (d) Box and whisker plots show 25-75 percentiles (box), the median (middle line) and 10/90 percentiles (whiskers). P value, non-paired Mann-Whitney test, 2 tailed. Data represent 43 cells each from 3 independent experiments. (e-g) Lack of expression regulation of DDRs, Syndecan 1 and CD44 in MV3 cells after stable β1/β3 knockdown. MV3 cells were cultured in 3D collagen matrix the absence or presence of mAb 4B4 and cRGD, harvested and analyzed by flow cytometry. (e) Expression of DDR1 and DDR2 after 9 h (f) Syndecan 1 and (g) CD44 after 24 h culture in 3D collagen culture. (e-g) Data represent 1 experiment. (h) Migration of MV3 β1/β3KD cells additionally treated with 4B4/cRGD and CD44 blocking mAb Hermes-1 compared with isotype control cells. Data obtained by tracking of 90 single cells for 24 h observation time. Data represent 1 experiment. Red lines, medians. P value, non-paired Mann-Whitney test, 2 tailed. See also **Figure 1**.

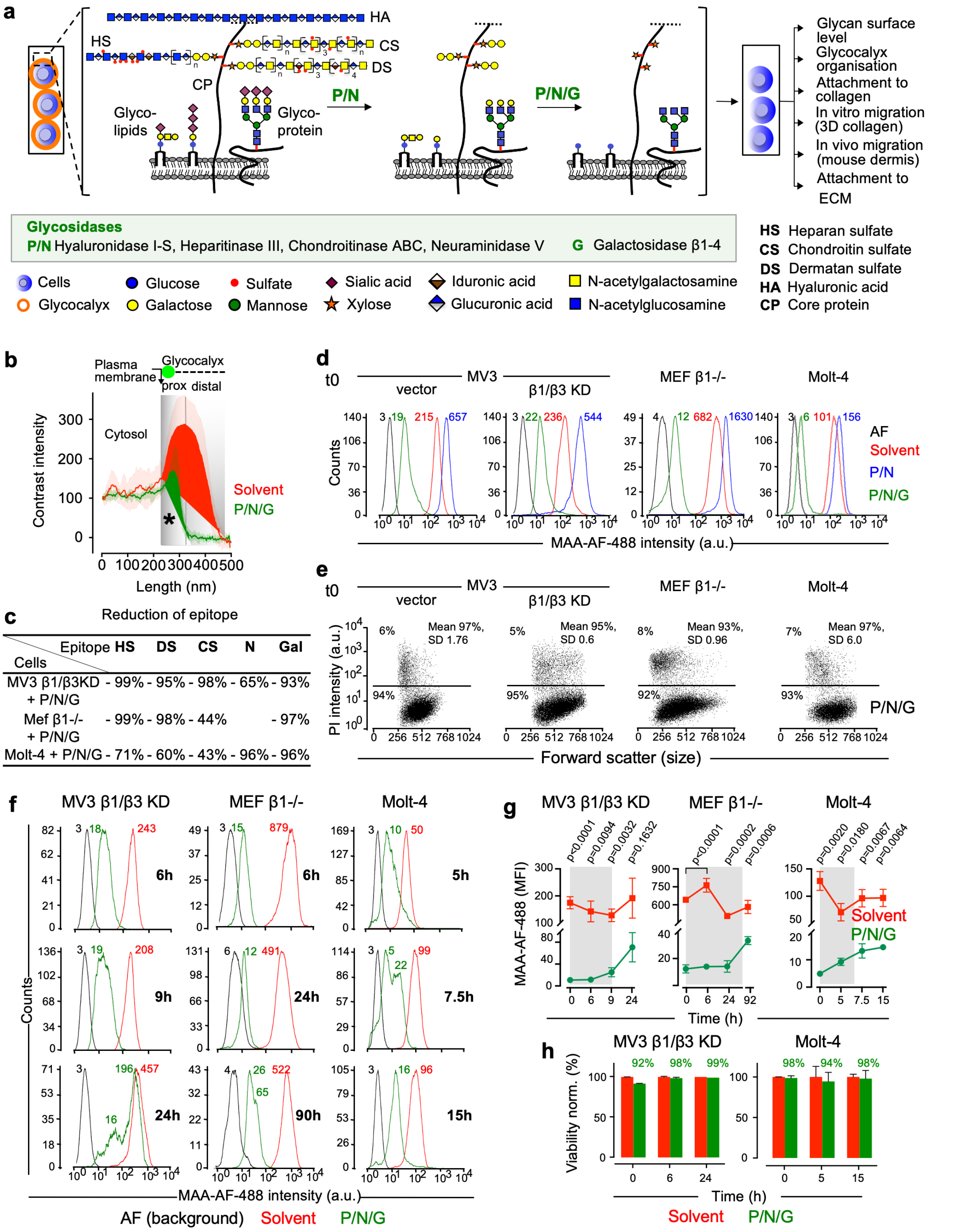


**Extended Data Fig. 4. Enzymatic removal of glycan moities on the cell surface of living cells using glycosidases and analysis of retention of the deglycosylated status.** (a) Schematic representation of cell surface glycocalyx structures, the stepwise enzymatic removal and the list of performed experiments. (b) Cumulative density of Ruthenium red staining before and after glycan removal resulting from 6 (MV3, solvent) and 4 (MV3, P/N/G) analyzed cells with 5 positions per cell of 2 independent experiments. Curves, averaged contrast intensity, shaded areas, SD. Blue dot, position of the plasma membrane. Dashed line, cross-sections perpendicular to the plasma membrane used for image analysis. Asterisk, interpolated gradient between intracellular and extracellular background signal used for calculating the integral. (c) Efficiency of the glycan removal using P/N/G analyzed by FACS and presented as reduction of the epitopes heparan, dermatan and chondroitin sulfate (HS, DS, CS) from 2 independent (MV3 β1/β3KD) or 1 experiment (Mef, Molt-4), sialic acids / neuraminic acids (N) from 1 experiment (MV3, Molt-4) and β1-4 Galactose (β1-4 Gal) from 5 (MV3) or 3 (Mef, Molt-4) independent experiments after treatment compared to untreated control cells. (d) Detection of glycan removal using fluorescently labeled Maackia amurensis agglutinin and (e) propidium iodide (PI) for cell viability in flow cytometry of MV3 integrin expressing and β1/β3KD and 4B4- and cRGD blocked cells, Mef β1-/- and Molt-4 cells. Mean (PI) viability results from 3 independent experiments. (f, g) Analysis of retention of the deglycosylated status of enzyme treated cells over a time span of 6-9 h (MV3), more than 90 h (Mef) and 5-6 h (Molt-4) and the stepwise recovery of β1-4 galactose expression. (d-f) 1 representative out of 5 (MV3) and 3 independent experiments (Mef and Molt-4). (g) Mean fluorescence intensities of Maackia amurensis agglutinin (MAA) with SEM of 50,000 cells from 5 (MV3) and 30,000 cells from 3 (Mef, Molt-4) independent experiments. (h) Determining the number of viable cells based on quantitation of the ATP presence of metabolically active cells. MV3 β1/β3KD and Molt-4 cells were untreated or treated with enzymes P/N/G for the 6 h and the viability was determined with the CellTiter-Glow® (CTG) assay at the indicated timepoints. The data shown is based on 150,000 cells per condition and experiment from at least 2 independent experiments (left panel, 0 h and 6 h), 1 experiment (left panel, 24 h) and at least 2 independent experiments (right panel). SEM is shown. P value, non-paired t-test, 2 tailed. See also **Figure 2 and 3**.

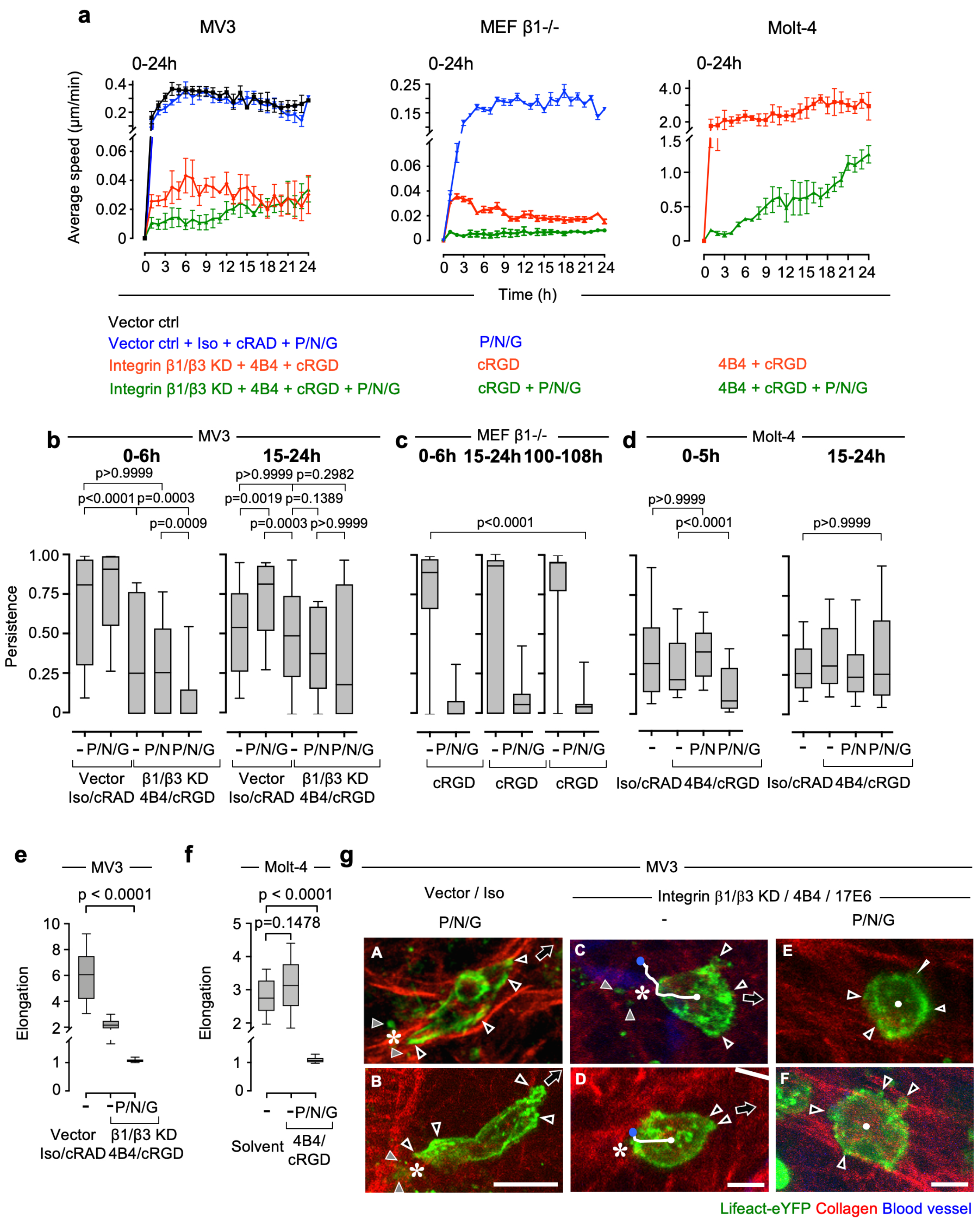


**Extended Data Fig. 5. Impact of glycan removal on migration, persistence and morphology of MV3, MEF and Molt-4 cells.** (a) Black line, average speed over 24 h of cells without integrin and glycan interference, (blue line) with glycan interference, (red line) after β1/3 integrin interference, and (green line) with additional glycan removal. Pooled mean average speeds with SEM of 120 (MV3), 90 (MEF) and 120 (Molt-4) cells from 3 independent experiments. (b-d) Impact of glycan removal on migration persistence of (b) MV3, (c) MEF and (d) Molt-4 cells. Data represent 87 (MV3, 6h), 111 (MV3, 15-24h), 51 cells (MEF) and 34 cells (Molt-4, 5h) and 43 cells

(Molt-4, 15-24h) for each condition from 3 independent experiments. (e, f) Impact of glycan removal on morphology of (e) MV3 and (f) Molt-4 cells. Elongation was determined as length/width of 99 cells for each condition embedded in 3D collagen I in 3 different experiments. (g) Re-onset of elongated phenotype of Lifeact-expressing MV3 vector control cells and amoeboid migration of MV3 β1/β3 integrin knockdown cells without and with surface glycan removal (P/N/G) after intradermal injection *in vivo*. ROIs of the imaging field of the dorsal skinfold chamber showing cells with typical morphological phenotypes within 3D collagen of the mouse deep dermis. White lines represent migration paths. Blue dots, starting points of migration. Arrows, direction of migration, based on retraction fibers and microparticles (g; A-C, grey-filled arrowheads) released from the cell rear (asterisks). Black-filled arrowheads, focal cell-collagen interaction sites (g; A, B), polarized (g; C, D) and diffused (g; E, F) distribution of actin-filled membrane blebs. Asterisks, cell rear. 2 representative cells for each condition from more than 10 cells analysed in 3 independent experiments. (g; A, B) Bar, 20 µm, (g: C-F) Bar, 10µm. (b-f) Box and whisker plots show 25-75 percentiles (box), the median (middle line) and 10/90 percentiles (whiskers). Black lines, medians. P values (b, d, e) non-paired Kruskal-Wallis with Dunn's post-test and (c, f) non-paired Mann-Whitney test, 2 tailed. See also **Figure 3**.

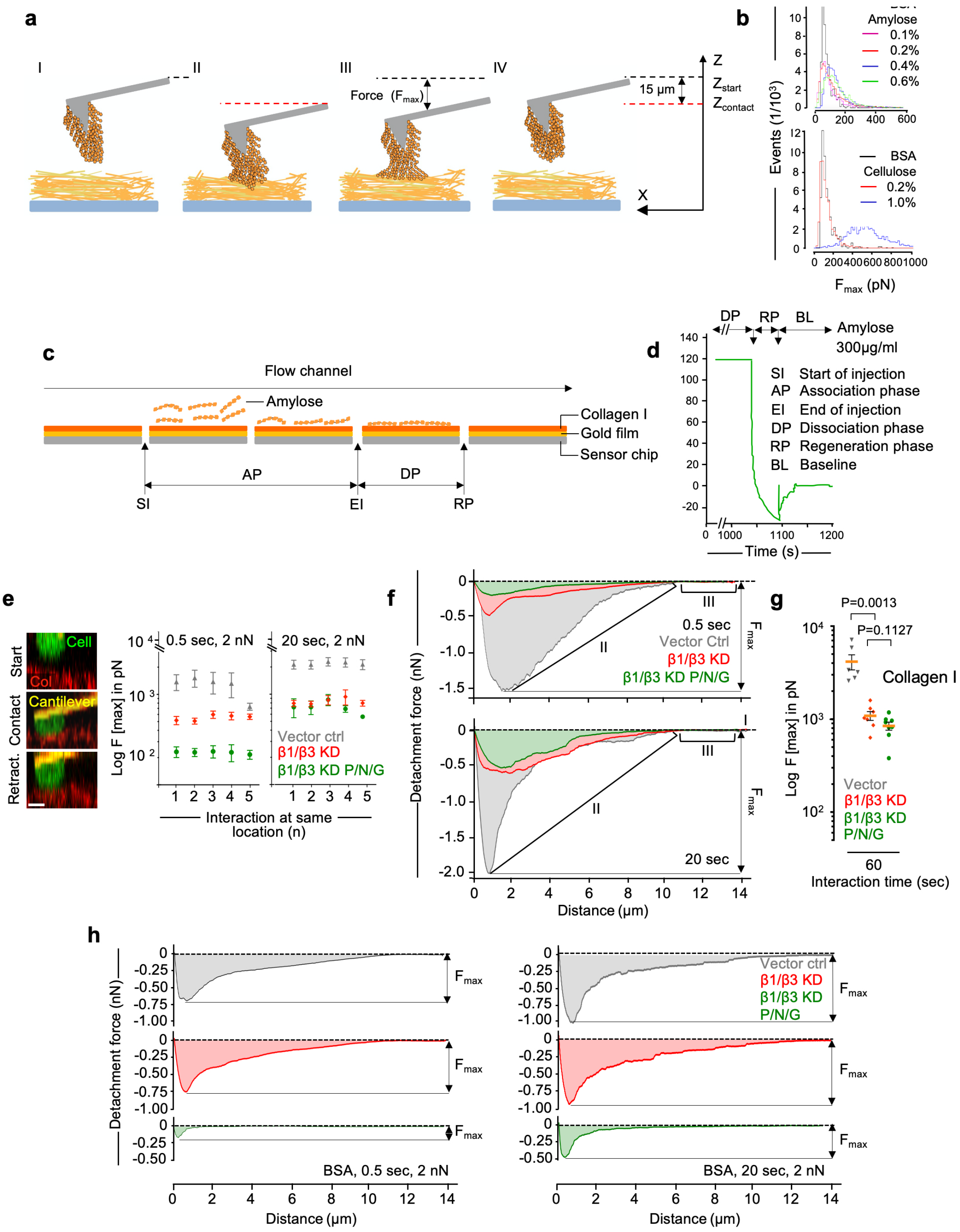


**Extended Data Fig. 6. Atomic force-based spectroscopy of glycan-binding to collagen fibers.** (a) Covalent coupling of multimeric polysaccharide chains to the surface of the cantilever tip to probe the force (F) required for cantilever detachment from a 3D fibrillar collagen surface. (b) Determination of F [max] with increased amylose and low- and high-concentrated cellulose coating of the cantilever, compared to background binding (BSA). (c, d) Glycan-mediated binding affinities to collagen fibers monitored by surface plasmon resonance (SPR). (c) Immobilization of bovine collagen I (orange layer) to a carboxymethyldextran-modified gold surface (yellow layer) of a CM5 sensor chip (grey layer) for biospecific interaction analysis in a surface plasmon resonance sensor. Amylose (orange squares) solution is injected as an analyte (SI) and binds to the stationary collagen I ligand during the association phase (AP) followed by the dissociation phase (DP) sustained by buffer flow and finally regeneration phase (RP). (d) Representation of amylose binding to collagen I after the regeneration phase (RP) obtained with a short pulse of 100 mM NaOH at the end of the experiment (wash-out), shown for the sample with the highest amylose concentration (300 µg/mL). 1 representative sensorgram out of 2 independent experiments. (f-i) Atomic force-based life cell spectroscopy of cell-surface glycan-binding to collagen fibers. (e) Coupling of a single cell to AFM cantilever to probe the force required for cell detachment F [max] from a 3D fibrillar collagen surface. Confocal microscopy of a single force measurement cycle between MV3 vector control and KD cells and collagen I surface, including start position, contact acquisition to the fiber and retraction. Life cell AFM single-cell force spectroscopy for 0.5 s and 20 s interaction time between MV3 cells and collagen I. Repeated cell – matrix interaction on the same collagen spot, showing similar magnitudes of generated forces at each binding event and equal force distribution between a cell and collagen (Mean and SEM, 9 cells, 3 independent experiments). Bar, 10 µm. (f) Averaged retraction curves of MV3 vector control cells (grey, 22 curves), β1/β3 integrin knockdown cells (red, 49 curves) and β1/β3 integrin knockdown cells with glycan removal (green, 50 curves) each of 11 cells from 3 independent experiments. (g) Interaction forces of MV3 cells to fibrillar collagen I after 60 s interaction time. Pooled values of maximum retraction forces to a mean value shown as Log F [max] and F [max] of 5 individual measurements per cell (orange lines show means with SEM) for MV3 vector control cells (grey, 6 cells), β1/β3 integrin knockdown without (red, 7 cells) or after glycan removal (green, 8 cells) on collagen per condition from 2 independent experiments. P values, non-paired t-test, 2 tailed. (h) Averaged control values for cell to bovine serum albumin (BSA) coated surface. MV3 vector control (0.5 s, 2 nN: grey, 28 curves; 20 s, 2 nN: grey, 42 curves from 9 cells), β1/β3 integrin knockdown cells (0.5 s, 2 nN: red, 45 curves; 20 s, 2 nN: red, 33 curves from 9 cells) and cells additionally treated with enzyme cocktail (P/N/G) (0.5 s, 2 nN: green, 34 curves; 20 s, 2 nN: green, 33 curves from 9 cells). See also **Figure 4**.

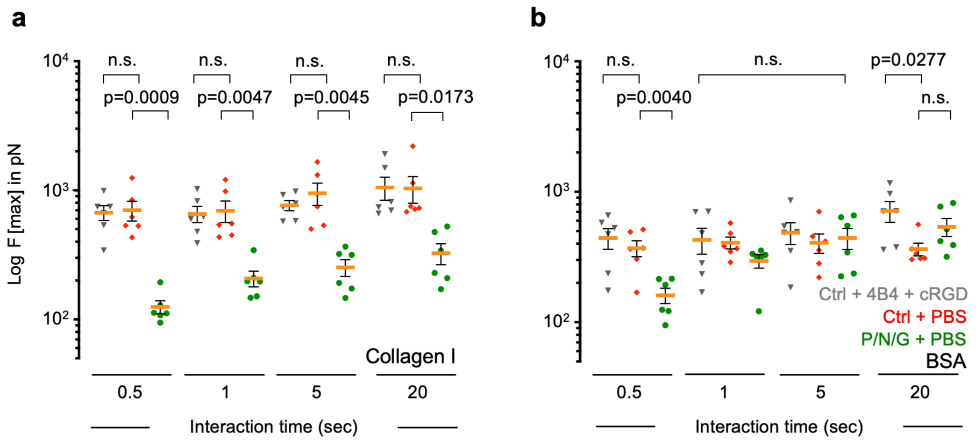


**Extended Data Fig. 7. Probing of glycan-binding to collagen fibers and studying the impact of enzymatic digestion of the glycocalyx on glycan-mediated cell binding to collagen. Atomic force spectroscopy of glycan-binding to collagen fibers.** (a, b) Live-cell atomic force spectroscopy of untreated Molt-4 control and additionally treated with P/N/G glycosidases. The maximum detachment force exerted on the cantilever F [max] was calculated from the peak to background level. Interaction forces of Molt-4 cells (6 cells per condition) to fibrillar collagen I (a) and BSA-coated surface (b) after 0.5 s, 1 s, 5 s or 20 s interaction time. Pooled values of maximum retraction forces to a mean value shown as Log F [max] of 3 individual measurements per cell (orange lines show means with SEM). P values (all graphs), non-paired t-test, 2 tailed. See also **Figure 4**.

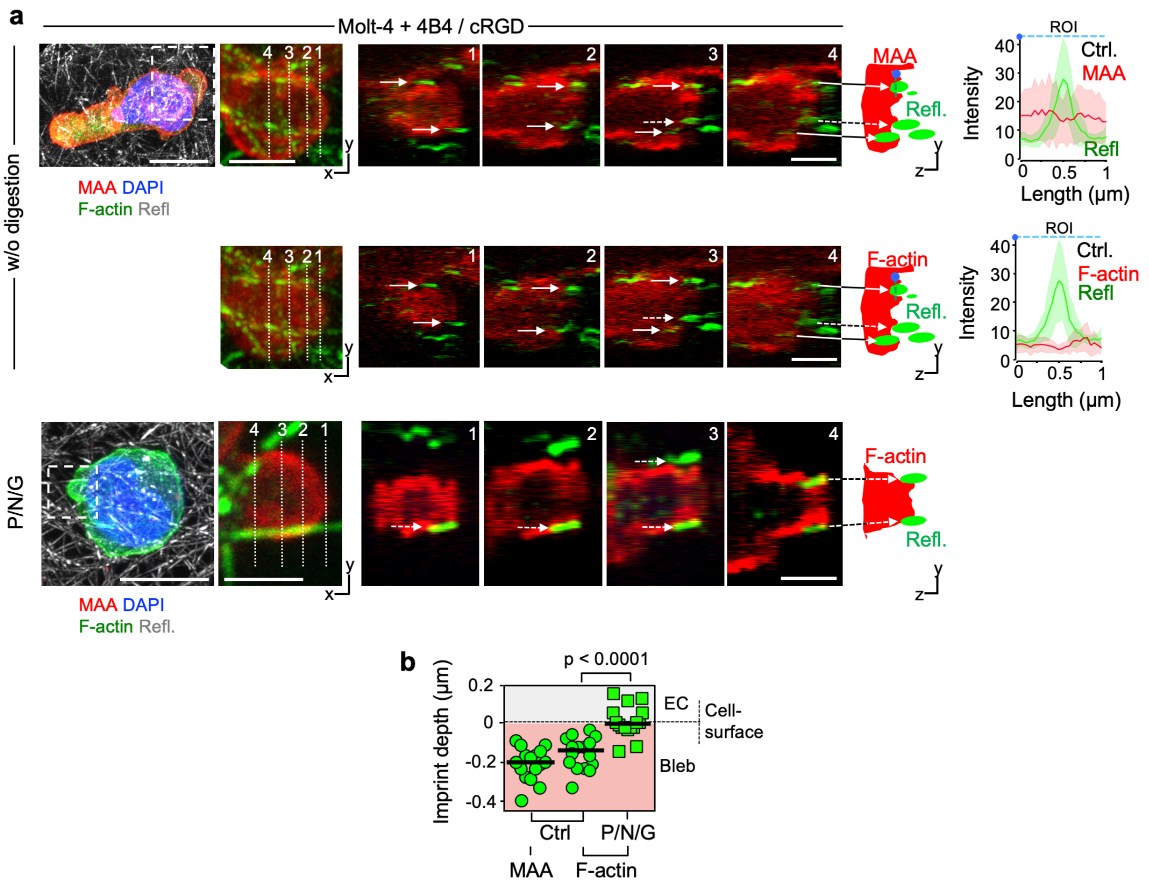


**Extended Data Fig. 8. Organization of glycan-dependent cell-matrix interactions.** Diffraction-limited confocal microscopy of glycan-dependent interactions with collagen fibers. Molt-4 cells were non-treated (a; upper panel, w/o digestion) or received P/N/G digestion (a; lower panel), embedded in 3D collagen and fixed after 90 min for detection of β-1,4 galactose residues (Maackia amurensis agglutinin, MAA), F-actin and collagen fibers (reflection). Images represent the xy and yz projections from serial z-scans as (dotted lines) with relative positions of cross-sectioned fibers (numbered 1-4) and cell surface represented as cartoons. Arrows, collagen fibers (green) located colocalized with MAA and F-actin-enriched protrusions (blebs, red). Dashed arrows, collagen fibers laterally intercalating with the cell surface. 1 representative cell out of 4 cells per condition analyzed in 2 independent experiments. Bars, 10 µm (overview), 2 µm (ROIs). Mean densitometry curves with SD of MAA and F-actin (red) and collagen fibril intensity (green) based on 20 curves in cross-sectioned cell blebs of 4 Molt-4 cells, analyzed in 2 independent experiments. (b) Median fiber imprint depth, based on MAA and F-actin signal from 16 curves in cross-sectioned cell blebs without and with glycan removal

(4 cells/ condition, 2 independent experiments). (b) Black lines, medians. P value, non-paired Mann-Whitney test, 2 tailed. See also **Figure 6**.

**Captions for Supplementary Videos**

**Supplementary Video 1.** Migration of β1+/+(fl/fl) and β1-/- murine embryonic fibroblasts (MEF) from multicellular spheroids in 3D collagen lattices (24 h, β1+/+(fl/fl), 48 h, β1-/- of observation).

**Supplementary Video 2.** Effects of β1/β3 integrin interference and enzymatic surface-glycan removal on the migration of MV3 melanoma cells in 3D collagen lattices. MV3 vector control cells in the presence of non-interfering isotype control antibody and cRAD peptide were compared with equally treated vector control cells after glycan removal using glycosidases P/N/G and MV3 β1/β3KD cells in the presence of mAb 4B4 and cRGD additionally incubated with solvent (PBS) or glycosidases P/N/G. Time-lapse sequences and path organization were monitored for 8h using bright-field microscopy. Time, hours:min. Bars, 20 µm.

**Supplementary Video 3.** Effects of β1/β3 integrin interference and enzymatic surface-glycan removal on the migration of MV3 melanoma cells in 3D collagen lattices. Extended time-lapse sequences and path organization of Movie 2 over 24h using bright-field microscopy. Time, hours:min. Bars, 20 µm.

**Supplementary Video 4.** Effects of enzymatic surface-glycan removal on the migration of β1 integrin-deficient murine embryonic fibroblasts in 3D collagen lattices. MEF β1-/- cells in the presence of cRAD or cRGD after treatment with solvent or glycosidases P/N/G were monitored for 24h using bright-field microscopy. Time, hours:min. Bars, 20 µm.

**Supplementary Video 5.** Effects of enzymatic surface-glycan removal on the migration of Molt-4 T lymphoma cells in 3D collagen lattices. Molt-4 cells incubated with 4B4 and cRGD after incubation in solvent or glycosidases P/N/G were monitored for 6 h using bright-field microscopy. Time, hours:min. Bars, 20 µm.

**Supplementary Video 6.** Relevance of surface glycans on MV3 cell migration *in vivo*. Lifeact-eYFP MV3 cells expressing β1/β3 integrin shRNA were additionally treated with solvent or glycosidases P/N/G, injected into the deep dermis of live nude mice and monitored for 6h by intravital multiphoton microscopy. Time, hours:min. Bars, 10 μm (Solvent), 20 μm (P/N/G).

**Supplementary Video 7.** Adaptation of migration mode and efficacy of MV3 melanoma cells after interference with integrins and removal of cell surface glycans. Cells were treated as described in Supplementary Video 2 and monitored by high-resolution bright-field microscopy for the indicated time periods (hours:min). Conversion from elongated to blebbing-amoeboid migration after β1/β3 integrin interference. Additional surface glycan removal caused immobilization despite ongoing bleb formation. Bars, 10 μm.

**Supplementary Video 8.** Inhibition of amoeboid migration of Molt-4 T lymphoma cells after removal of surface glycans. Molt-4 cells were treated as described in Supplementary Video 4 and monitored by high-resolution bright-field microscopy for 3.5 h. Surface glycan removal caused transient immobilization despite ongoing cytoskeletal dynamics (“running on the spot”). Time, hours:min. Bars, 10 μm.
